# Structure of a contractile injection system in *Salmonella enterica* subsp. *salamae*

**DOI:** 10.1101/2025.06.01.657177

**Authors:** Rooshanie N. Ejaz, Kristin Funke, Claudia S. Kielkopf, Freddie J. O. Martin, Marta Siborová, Ivo A. Hendriks, Nicholas H. Sofos, Tillmann H. Pape, Eva M. Steiner-Rebrova, Michael L. Nielsen, Marc Erhardt, Nicholas M. I. Taylor

## Abstract

Extracellular contractile injection systems (eCISs) are phage-derived nanomachines used by bacteria to deliver effectors into target cells. Well-studied examples include the *Photorhabdus asymbiotica* virulence cassettes and the antifeeding prophage from *Serratia entomophila*, which have been engineered for heterologous cargo delivery. Recent genomic analyses identified previously uncharacterized eCIS gene clusters in the opportunistic human pathogen *Salmonella enterica* subspecies *salamae*, but their structure, function, and biotechnological potential remain unexplored. Here, we report a high-resolution cryo-electron microscopy structure of the *S. enterica* eCIS. Our atomic models reveal a unique sheath architecture, an expansive cage-like shell around a central spike, and an associated transmembrane hydrolase. We identify a putative effector encoded within the operon exhibiting periplasmic toxicity and provide evidence that the *S. enterica* eCIS deviates from canonical eCISs by interacting with the inner membrane. Guided by these structural features, we uncover a previously unannotated cluster of contractile injection systems (CISs). Together, our findings expand the known diversity of CISs’ structures and functions and lay the groundwork for engineering customizable protein delivery platforms.

## Main

Extracellular contractile injection systems (eCISs) are macromolecular complexes expressed by bacteria and archaea^1,2,3,4,5^. Evolutionarily derived from the contractile tails of bacteriophages^6,7,8^, they assemble into nanoscale injectors^6^, capable of targeted piercing of prokaryotic and eukaryotic cell membranes^5,9,10^ to deliver protein payloads. Bacterial cells synthesize eCISs and presumably undergo lysis upon a triggering event^2,11^, allowing these systems to leave the cytoplasm and inject effectors across membranes of target cells^12,13^. eCISs have been bioengineered to deliver customized payloads as therapeutics and biopesticides^12,14^. Studying eCIS function and structure in human pathogens, such as *Salmonella* species, could provide biomedically relevant insights into novel injection systems.

The genome-wide prediction of 631 eCIS-like co-translational assemblies by Chen *et. al*. revealed a complex genetic landscape of eCISs^1^, identifying several eCIS homologues in the genus *Salmonella*^1^. *Salmonella enteric*a, a Gram-negative, flagellated, rod-shaped enterobacterium^15^, includes subspecies *salamae* and *diarizonae,* both of which encode eCIS-like clusters, of which 143 were identified in *salamae*^1^. These two subspecies are typically isolated from environmental sources, such as soil samples, and occasionally reptilian guts^16,17^. *S. salamae* has been known to cause opportunistic infections in immunocompromised patients^18^. Notably, *salamae* subspecies have evolved to survive outside of their host, compared to pathogenic serovars^17^.

Canonical eCISs comprise a baseplate, long tail fibres, and a contractile sheath surrounding an inner tube terminating in a cap^1,5,6,9^. The most well-studied of these canonical extracellular systems are the Antifeeding prophage (Afp)^9,19,20^ and the *Photorhabdus* virulence cassettes (PVC)^5,12,13^. eCISs share structural features with contractile injection systems (CIS), such as Type VI secretion systems (T6SS), including their outer membrane anchor deficient subtypes, (T6SS^ii-iv^) and tailocins^6,21^. AlgoCIS, tCIS and CIS^Sc^ from *Algoriphagus machipongonensis,* cyanobacterium *Anabaena* and *Streptomyces coelicolor*, respectively, were structurally characterized to help elucidate their diverse functions^3,4,22^. CISs can be considered interactive protein complexes, allowing trans-kingdom interactions and involvement in bacterial cell cycles. The role of CISs as interaction tools across bacterial species and clades is expanding^2,3,22^.

In this study, we solve the first atomic structure of a CIS in *Salmonella*, including its membrane-associated component. We then apply structural analysis to demonstrate how the *Salmonella* CISs (SalCIS) may deviate from canonical eCISs by anchoring to the inner membrane of *S. enterica*. We leverage structural insights to identify a previously unannotated and uncharacterised cluster of bacterial CISs. Finally, we characterise two putative effectors present in the operon, one of which exhibits periplasmic toxicity. Understanding these mechanisms may provide deeper insights into the evolutionary and functional diversity of eCISs in pathogenic bacteria.

## Results and discussion

### SalCIS is a novel contractile injection system with conserved CIS architecture

The *Salmonella enterica* subspecies *salamae* genetic cassette is a co-translated bidirectional operon, comprising eighteen open reading frames (Fig. 1a)^1^. Bioinformatic analysis identified four strains of *Salmonella enterica* subsp. *salamae* encoding a CIS cassette^1^ (SalCIS). Out of these previously uncharacterised operons, the operon from strain NCTC9936 was selected for further study, due to continuous translation sequences. The Afp nomenclature (Afp1 to Afp16) was used to assign a putative function to all hypothetical proteins (SalCis1-21). The putative packing AAA+ATPase SalCis15 is encoded upstream of the putative effector proteins SalCis20 and SalCis21 (Fig. 1a). According to previous eCIS classification^1^, SalCIS falls into clade Ib, which includes AlgoCIS from the marine bacterium *Algoriphagus machipongonensis* and the T6SS subtype IV (T6SS^iv^) from *Asiaticus amoebophilus* (Fig. 1a). Notably, a putative protein, SalCis19, was predicted to have transmembrane helices, and a peptidoglycan hydrolase domain, with conserved homology (Extended data Fig.1a-c). To understand the SalCIS system via structural analysis, a deletion mutant of the SalCIS operon (ΔSalCis13-ΔSalCis19) was cloned, heterologously expressed in *E. coli* and purified for cryo-electron microscopy (cryo-EM). Maps at resolutions of 2.86 Å (baseplate), 2.3 Å (extended sheath) and 2.97 Å (cap) were obtained (Fig.1c-f). The density was excellent throughout, apart from a tapering off in resolution towards the edges of the baseplate maps. Though detected in LC-MS data (Fig. 1g and Supplementary Table 1), SalCis15 was absent from all maps, in line with a transient association with SalCIS, as noted in other CIS^10,19^.

**Figure 1:**
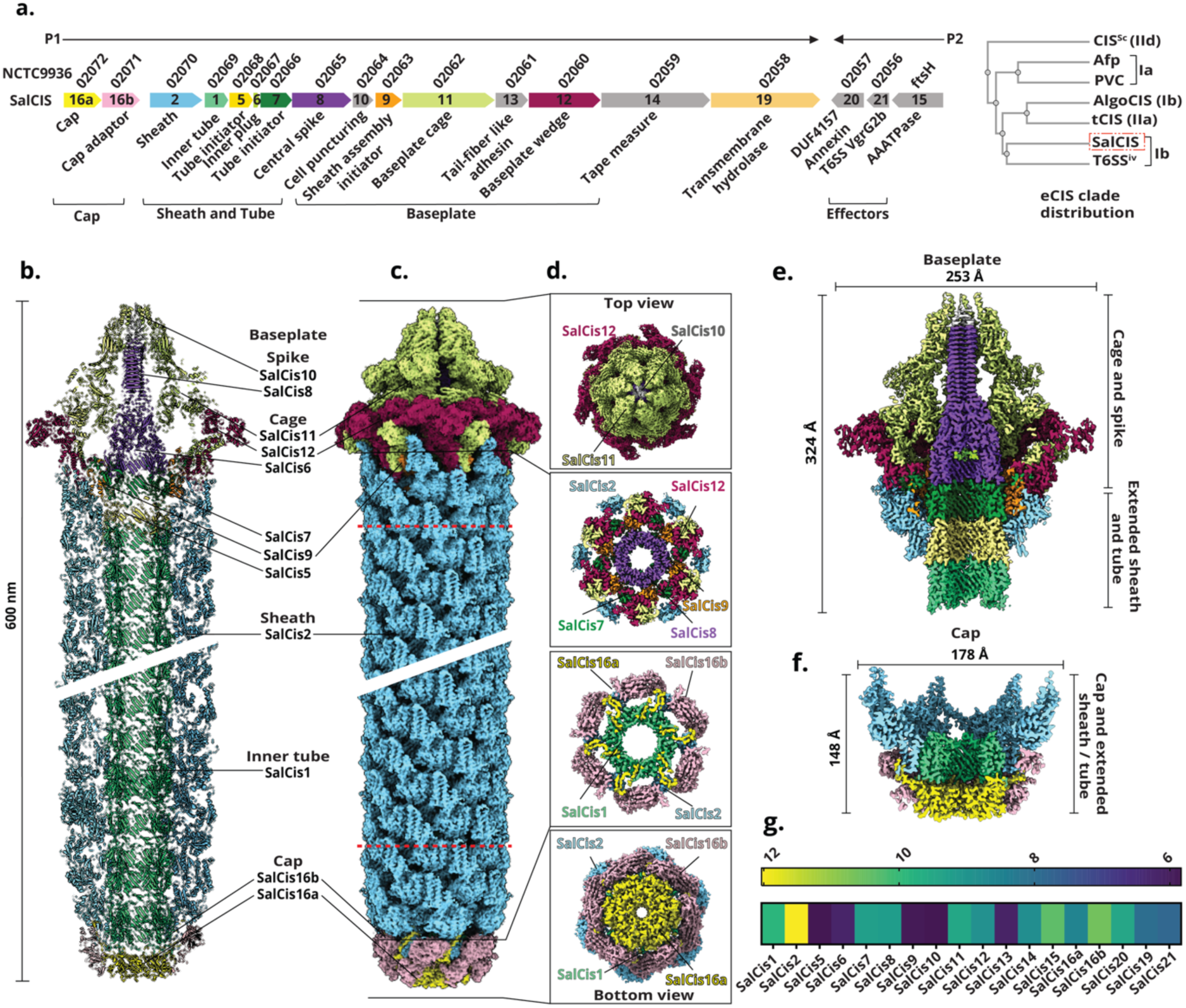
Atomic structure of the *Salmonella* CIS. **a:** *Salmonella* CIS (SalCIS) cassette, with genes annotated according to the NCBI genomic database identifiers, and numbered according to the antifeeding prophage nomenclature. Descriptors of genetic components are presented according to resolved densities, all coloured components were fully or partially resolved in cryo-EM maps, grey components were not. The text under the brackets indicates assemblies and/or functions confirmed in this study. P1 refers to the first natural promoter identified, and P2 refers to the second one. Right panel shows placement of SalCIS within eCIS clade organization. **b:** Atomic structure of SalCIS as cartoon representation, clipped at the centre vertically to reveal tube and central spike features. Model depiction is a stitched representation of symmetry copies of constructed atomic models expanded onto the cryo-EM structure. **c:** SalCIS cryo-EM density. The map shown is a composite map constructed by merging three separate maps, dashed red lines indicate borders where maps were stitched, σ for each map is reported in Supplementary Fig. 7 and 8. **d:** Boxed lateral insets show cross-sectional slice views illustrating the arrangement and interactions of various SalCIS proteins within the baseplate and cap complexes. **e:** Clipped view of the SalCIS baseplate complex cryo-EM map, highlighting individual units and interactions between the baseplate components, comprising the outer cage (SalCis11 and SalCis12) and the inner spike and puncture complex with the tube and sheath vertex (SalCis10, SalCis8, SalCis7, SalCis5, SalCis2 and SalCis1). **f:** Clipped view of the SalCIS cap complex cryo-EM map highlighting individual units and interactions of the terminal cap complex with the inner tube (SalCis16a, SalCis16b, SalCis2 and SalCis1). **g:** LC-MS coverage of SalCIS proteins, heat map shows log^2^ of unique spectral counts of all proteins present, reported in Supplementary Table 1, coloured according to key.

SalCIS is composed of a baseplate, an extended sheath tube complex, and a terminal cap (Fig.1a). The baseplate surrounds the central spike in a cage and compact wedge complex, that transitions into the sheath and tube (Fig. 1e, insets). The extended sheath and tube complex assembly interacts with the cap (Fig. 1f). The novel architecture of this CIS preserves key features noted in other CISs, such as bacteriophages, T6SS, and previously described eCISs (Supplementary Table 2).

The SalCIS inner spike retains the structure of eCISs such as AlgoCIS and tCIS, with a SalCis8 trimer, and capped by a single PAAR-like SalCis10 at its C-terminus (Fig. 2a)^23^. The gp27- and gp5-like ladder connected through a β-barrel are also retained (Extended Data Fig. 2a and b). The gp5-like ladder of SalCis8 is about 27 Å longer than eCISs like PVC and Afp and instead resembles the spike arrangement of clade Ib eCISs (Fig. 2b). The gp27-like domain of one SalCis8 monomer interacts with the proximal end of two copies of SalCis7 primarily through electrostatic interactions, via defined residues. The LysM domains of SalCis7 stabilizes the spike through contacts with SalCis8 (Extended Data Fig. 2c). The putative plug protein, SalCis6, encoded with an alternative start codon (Val), is inferred by LC-MS data (supplementary table 1.). Its complete density could not be resolved, due to particle scarcity, only α-carbon chain loops that correspond to the residues 49-56 of the AlphaFold3 model were traced within the lumen (Fig. 2c), indicative of a conserved spike plugging mechanism in SalCIS^4^.

**Figure 2:**
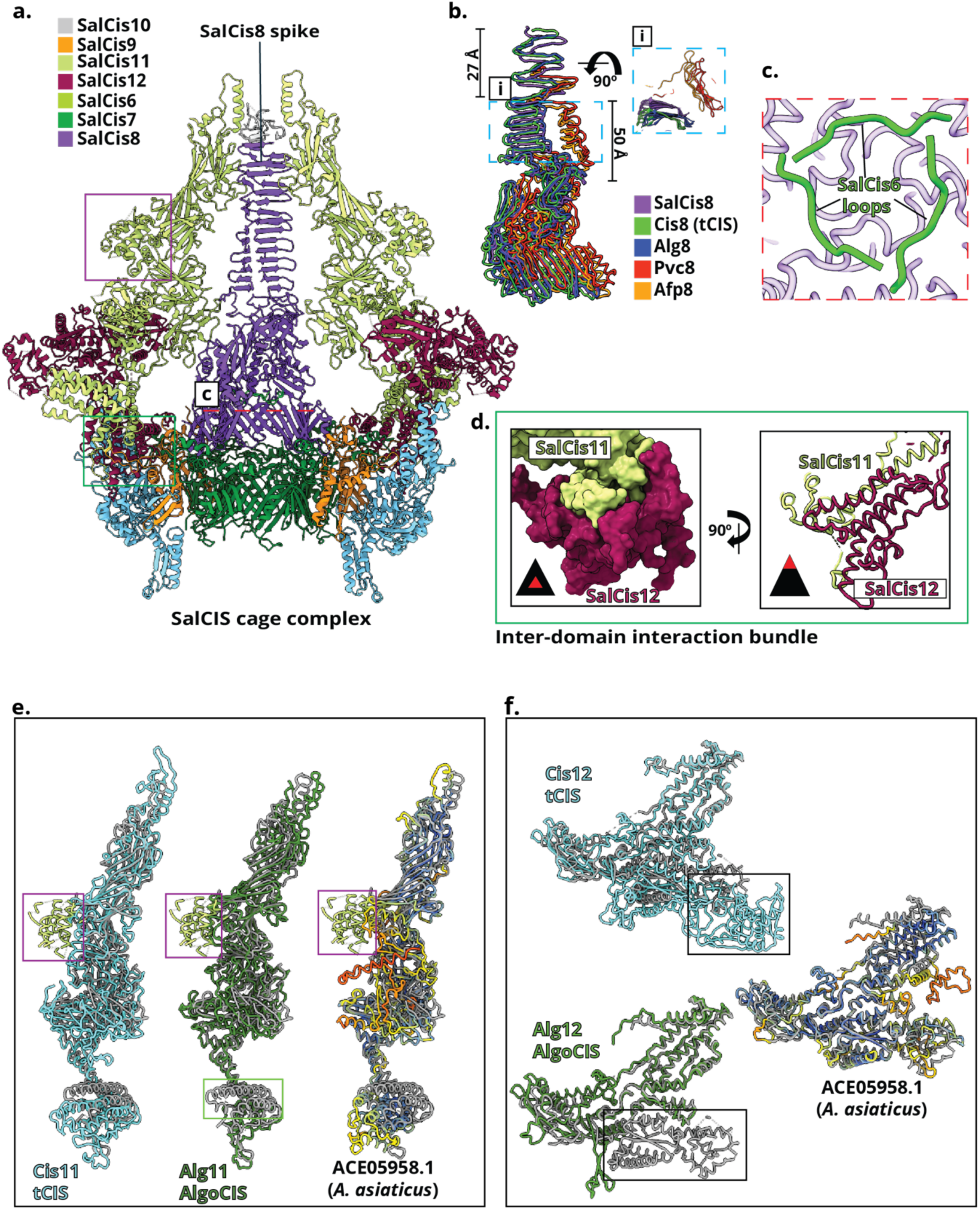
Details of the spike-puncture complex. **a:** Cartoon depiction of SalCIS inner spike and cage complex, chains are coloured according to key. Surrounding chains are hidden to reveal interaction details. Red dashed line highlights features detailed in inset c. Green box highlights features detailed in inset d. Purple box highlights feature detailed in e. **b:** Cartoon depiction of the SalCis8 spike structure, highlighting conserved domains. Blue dashed box shows top-down view of the spike "i.". Various spike proteins are coloured by key. **c:** SalCis6 viewed within the inner lumen of SalCis8, from perspective of red dashed line in a. Depicting three loops traced within the spike density, corresponding to the AlphaFold3 model of SalCis6. **d:** Shows domains I and IV from SalCis12 interacting with SalCis11’s domain I via α-helices. Left panel depicts electrostatic surface depiction of the solved structure, right panel depicts side view of the same in chain format. **e:** SalCis11 has a unique extension, as shown by superimposition of the solved structure of SalCis11(in grey) onto homologues, purple boxes indicate an extension of the cage (coloured green), only present in SalCis11. Green box indicates helical extension of SalCis11’s domain missing in AlgoCIS. **f:** SalCis12 is related to other CISs as illustrated by the superimposition of the solved structure of SalCis12 (in grey) onto homologues. Black box around Cis12 from tCIS indicates large domain extension not present in SalCis12, black box around SalCis12 extended domain indicates extension of domain in SalCis12, not present in AlgoCis12. The predicted structure of the SalCis12 homologue in *A. asiaticus* highlights high structural similarity. The *A. asiaticus* cage protein homologues are AlphaFold-predicted structures coloured by pLDDT confidence scores, where blue indicates high confidence and orange/red indicates low confidence.

The spike assembly is surrounded by a cage, including hexameric assemblies of both SalCis11 and SalCis12, forming a conserved trifurcation unit akin to the baseplate of the bacteriophage T4 and other eCISs (Extended Data Fig. 3a)^6,8^. SalCis11 and SalCis12 display gp6 and gp6/7-like folds, respectively^6,8^ (Extended Data Fig. 3b-d). SalCis11’s domain I extensively interfaces SalCis12’s domains I and IV via α-helical contacts (Fig. 2d). Notably, SalCis11 harbours a unique, extended domain IV that was not identified in other CISs (Fig. 2e). Surface analysis of the seven α-helices (residues 419–549) comprising this domain, revealed a distinct charge distribution, coupled with a hydrophilic character (Extended Data Fig. 3e).

SalCIS12 is deficient in a large extension of domain V, as is present in Cis12 from tCIS, and extended in comparison to its homologue Alg12 from AlgoCIS (Fig. 2f), resulting in a compact baseplate wedge, nearly hexagonal in shape in SalCIS (Extended Data Fig. 3a). SalCis12 is instead homologous to the predicted structure of the putative cage proteins of *A. asiaticus*, (Fig. 2f). In *A. asiaticus* T6SS^iv^, injection systems form bundles^21^. We noted the presence of CIS aggregates in our TEM analysis (Supplementary Fig. 1). It is plausible that the compact nature of the *Salmonella* CIS baseplate facilitates the occurrence of these injection systems in bundles wherein the baseplate is tethered to the inner membrane. The outer surface of the baseplate is dominated by positive charges, around the circular tip formed by the extended domain IV (Extended Data Fig. 3e). The distal tip is also positively charged. The combination of positive charge with hydrophilic character likely facilitates membrane penetration^3,20,24^. Taken together, our analysis indicates that SalCIS’s baseplate may be a membrane-interacting variant. It combines conserved eCIS architecture with a more compact geometry, forming bundles. Electrostatic charges around the baseplate are favourable for penetrating a membrane, with conserved homology closer to clade Ib eCISs.

### SalCIS is composed of a long sheath and tube vertex

SalCis2, the sole sheath component of SalCIS is divided into four domains (Fig. 3a). An X-loop domain was resolved, not previously noted in eCISs. The X-loop forms salt bridges between ASP322–ARG84 (P3) and LYS43 (P4), of two sheath protomers (Fig. 3b, i.), stabilising the extended conformation. Domain III includes two α-helices and a disordered loop, distinctly oriented flush parallel with the sheath wall (Extended Data Fig. 4 a, i.).

**Figure 3.**
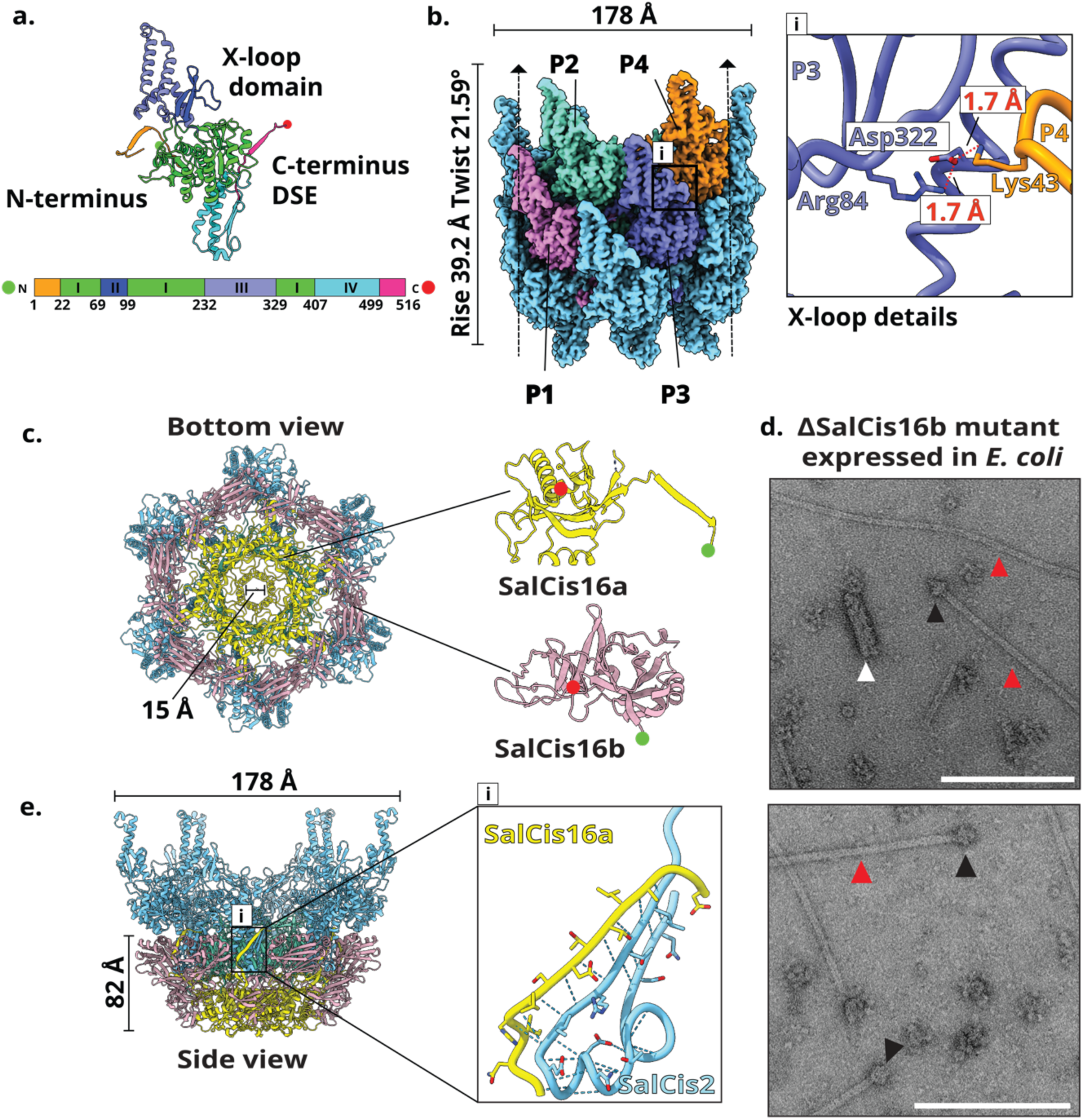
Outer sheath and tube terminate at the cap. **a:** Domain organization of SalCis2. **b:** Details of outer sheath assembly. Four protomers (P1-P4) are highlighted and color-coded; black box indicates a zoom-in of key inter-protomer contacts, with corresponding cryo-EM density shown. Vertical dashed lines indicate vertex wall, arrowheads point towards baseplate. (i) Shows details of the X-loop motif, key residues are highlighted, and atomic distances marked. **c:** Hexameric structure of the SalCIS cap shown in cartoon format, viewed from the bottom. N- and C-termini (red and green, respectively) of SalCis16a and SalCis16b monomers are indicated, the structures shown in cartoon format. **d:** Negative-stain TEM images of a ΔSalCis16b mutant expressed in *E. coli,* purified by gradient fractionation. Black triangles indicate baseplate regions of tube–baseplate complexes; red arrows mark polymerized inner tube complexes; white arrow shows low-energy sheath assemblies. Scale bar, 200 nm. **e:** Cartoon depiction of the SalCIS cap structure. Coloured box indicates zoomed-in views of key contact sites. (i) Top box shows a model of structural interactions between the final DSE motif between SalCis2 tube and SalCis16a, polar residues and hydrogen bonds are highlighted

The sheath adopts a helical symmetry, with a rise of 39.2 Å and a twist of 21.6°. Sheath assembly is stabilised by a conserved C-terminal donor strand exchange (DSE) motif. The N-terminus of each SalCis2 monomer (P3) engages in a tripartite DSE motif, forming between protomers 1 and 2, previously noted in tCIS (Extended Data Fig. 4a, ii and iii.).

Three distinct conformational states of SalCis2 are observed (Extended Data Fig. 4b). The C-terminus of R1 shifts (residues 507–516) to accommodate DSE with SalCis9, distinct from R2/R3. The R1 N-terminus also shifts to connect with the helical SalCis2-R2 (Extended Data Fig. 4b). Comparative analysis revealed SalCIS shares distant homology with Alg2, Cis2 (tCIS), and *Streptomyces* Cis2 (Extended Data Fig. 4c). The AlphaFold3 structural prediction of the *A. asiaticus* sheath protein (ACE06419.1) is predicted to harbour the X-loop, but it does not retain the vertical helix-loop-helix motif orientation (Extended Data Fig. 4d). Structural homology searches revealed a unique cluster of hits, with single sheath protein CISs, structurally identical to SalCis2 (Extended Data Fig. 5a and b).

Cryo-EM reconstruction of the contracted sheath informed an estimate of 272 Å length (Extended Data Fig. 6a and b). This estimate would result in moving the inner tube forward by a distance of ∼328 nm. Sufficient for piercing *S. enterica salmae’s* own double membranes^25^ (Supplementary formula 1).

SalCIS1 is highly conserved; superimposition of SalCIS1 with Alg1, Afp1, Cis1 from *Streptomyces*, Pvc1, and Cis1 reveals strong structural similarity. So does comparison of SalCIS1 with the structure prediction of the *A. asiaticus* Hcp component (Extended Data Fig. 7a-c).

Taken together, our analysis of the tube and sheath complex shows that the inner tube of CIS’s including SalCIS, seems to be highly conserved, but the tail sheath of SalCIS appears to have acquired some unique adaptations that might have important functional roles.

### SalCis16b is the cap adaptor of SalCIS sheath vertex

The first gene in the *S. enterica* SalCIS cluster, SalCis16a, forms a hexameric cap structure with an opening of 15 Å between the distal-most helical loops (residues 143-155). This is wider than the opening reported in the distal loops of the single cap-protein assembly of the Afp cap, belonging to clade Ia eCIS^20^. In contrast to clade Ia eCIS, a second hexamer forms the complete cap in SalCIS, via the terminal cap adaptor, SalCis16b (Fig. 3c). Previously, a double cap assembly was reported in the AlgoCIS (clade Ib) and tCIS (IIa), with similar dimensions^3,4^. Deletion of SalCis16b, resulted in disrupted particle assembly, and the formation of tube– baseplate complexes (Fig. 3d). Negative-stain TEM revealed low-energy assemblies of the sheath protein SalCis2, with no extended or contracted sheath conformations assembled on the baseplate (Fig. 3d). The final DSE interaction, between SalCis2 and the cap, is stabilized by the N-terminus of SalCis16b, the cap adaptor. Here, the SalCis2-R3 conformation, engages in a terminal DSE motif with SalCis16a (Fig. 3e). The N-terminus of SalCis1’s final six protomers show a distinct change in conformation, upon binding Sal16a (protomer 2, Extended Data Fig. 7d). Similar differences in conformation of SalCis1 homologues has been reported in eCISs such as the Afp^20^.

Together, these interactions define an ordered cap assembly, where SalCis16b anchors SalCis2 via the terminal DSE, closing the sheath-tube complex

### SalCis19 is an integral membrane protein containing a putative hydrolase domain

SalCis19 was predicted to have three transmembrane α-helices and a Sec-type signal sequence with a cleavage site at residue 42 (Fig. 4a)^26^. Cryo-EM analysis of LMNG-solubilized SalCis19 resulted in a 2.73 Å resolution map of the homodimer. Each monomer contains three transmembrane α-helices, embedded in the detergent micelle, and a large periplasmic domain (Fig. 4b and Supplementary Table 1, Supplementary Fig. 2). Residues 1-42 were absent in the mass spectrometry analysis of the purified protein, which may indicate post Sec-mediated export cleavage^27^ (Fig. 4a). The postulated periplasmic domain adopts a conserved hydrolase fold, homologous to SleL (PDB:4s3j), a cortical spore lytic enzyme from *Bacillus cereus*^28^ (Fig. 4c), with conservation of several residues in the catalytic core (Fig. 4d, RMSD 0.976 Å). SleL is an N-acetylmuramoyl-L-alanine amidase, which cleaves peptidoglycan^28^. Given the similarity of *Salmonella* and *Bacillus* peptidoglycan composition^29^, we propose that SalCis19 remodels the cell wall, to assist SalCIS insertion, analogous to MltE insertion by *E. coli* T6SS^30^. Inspected genomes of CIS clade Ib and structure-based searches^31^ revealed putative homologues of SalCis19, with similar signal sequences, topologies and domains (Extended Data Fig. 1a and b)^32,33,34^.

**Figure 4.**
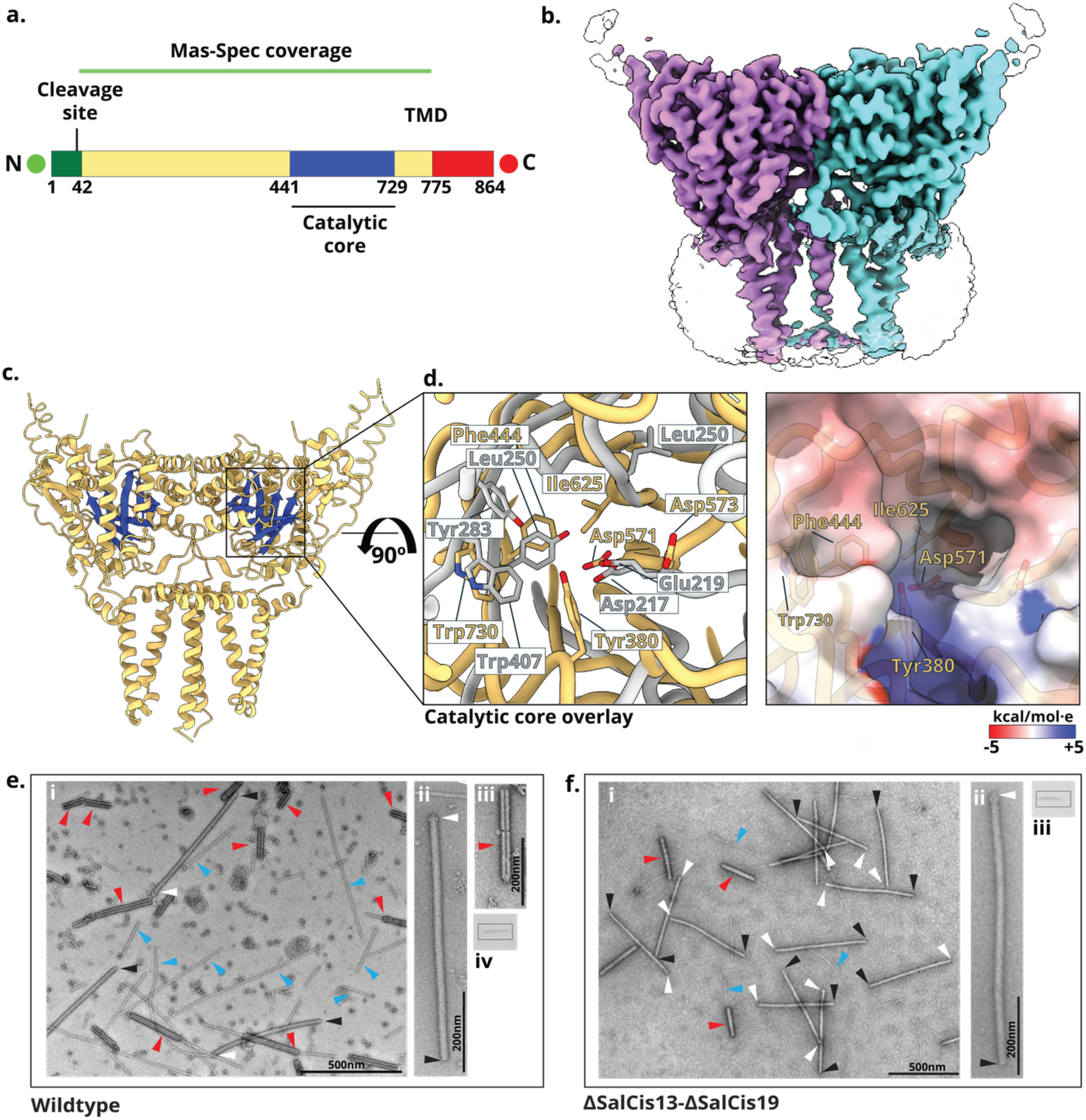
Cryo-EM reveals the structure of an integral membrane protein in SalCIS. **a:** Predicted domain organization of SalCis19. Bioinformatic analysis shows an N-terminal Sec-type signal sequence (cleavage after residue 42), three TM α-helices per monomer, and a large periplasmic domain (residues 44–775), green bar indicates mass-spec coverage, transmembrane domain is indicated (TMD). **b:** Cryo-EM structure of LMNG-solubilized SalCis19, σ 3.61. Reconstruction reveals a homodimer, with transmembrane (TM) helices embedded in a detergent micelle and a large periplasmic domain extending into the solvent, each monomer coloured. **c:** Solved structure of SalCis19 homodimer, highlighting the two catalytic cores in blue. **d:** Left panel shows structural alignment of the catalytic cores from *B. cereus* SleL (grey) and *S. enterica* SalCis19 (cantaloupe), highlighting conservation within the catalytic domain. Right panel shows negative electrostatic surface charge localised in the putative substrate binding core. **e:** Negative stain images of the wildtype SalCIS operon expressed and purified from *E. coli.* Black arrowheads point at caps, white arrows point at baseplates, red arrows denote contracted sheath assemblies, blue arrows denote expelled inner tubes. (i) Low magnification image of diluted final preparation of heterologously expressed wild-type SalCIS in *E. coli* (BL21 DE3). (ii) Magnified view of one extended SalCIS particle. (iii) Magnified view of contracted sheath assembly. **f:** Negative stain images of ΔSalCis13-ΔSalCis19 mutant of SalCIS expressed and purified from *E. coli* (BL21 DE3). Labelled the same as e. (i) Low magnification image of diluted final preparation. (ii) Magnified view of one extended SalCIS particle. Western blot band indicates presence of SalCis2 at 57 kDa in e(iv) and f(iii), probed with an anti-SalCis2 specific antibody, cropped for clarity.

### Structural modelling suggests SalCis13 acts as a membrane-associated anchor

SalCis13 is the putative tail fibre of SalCIS. Structure-based searches revealed highly conserved homologues (Extended Data Fig. 8a). AlphaFold3 modelling of SalCis13 as a trimer was more favourable than other oligomeric states tested, distinct from canonical eCIS tail fibres (Extended Data Fig. 8b). Structural similarity was noted when compared to the AlphaFold3 structure of *A. asiaticus* T6SS^iv21^ T6SS^iv21^ (Extended Data Fig. 8b). The baseplate of SalCIS was docked into the sub-tomogram averaged structure of the *A. asiaticus* CIS, revealing similar geometry (Extended Data Fig. 8c). Modelling of SalCis13, SalCis19, and SalCis11 as a complex located the C-terminal TMD’s of SalCis19 engaging the trimer of SalCis13, which in turn binds SalCis11 via the N-terminus of SalCis13 (Extended Data Fig. 8d).

Expression of the full SalCIS operon in *E. coli* and *S. enterica* yielded mostly aggregated or disassembled particles, unsuitable for structural analysis (Fig. 4e and Supplementary Fig. 1 and 3a). LC-MS analysis detected SalCis19 post purification at a relative abundance of 0.005, normalized to the cap protein SalCis16b at 1.0, (Supplementary Table 1.). The low abundance may indicate dissociation during lysis, reminiscent of the membrane associated complex in T6SSs^35^. Deletion of SalCis13 and SalCis19 individually did not improve particle yield or quality (Supplementary Fig. 3b). The double deletion mutant (ΔSalCis13-ΔSalCis19) yielded the highest proportion of intact particles and enabled high-resolution single particle reconstruction (Fig. 4f).

Taken together, our analysis of SalCis13 and SalCis19 was used to guide a hypothesis of how SalCIS may tether to the inner membrane (Extended Data Fig. 8d)^36^.

### The SalCIS expresses effectors toxic to bacteria

Within the SalCIS operon, we identified two putative effectors, positioned near the AAA+ATPase, SalCis15. For LC-MS analysis, the SalCIS operon was co-expressed in *E. coli*. SalCis1-19 were expressed on a pBAD33 plasmid, whereas SalCis 15, 20 and 21 were expressed on a pET11a plasmid. After purification, both putative effectors were detected by LC-MS in a gradient-separated preparation, together with the core components of SalCIS, consistent with their retention, at least in some of the particles, during complex isolation (Supplementary Table 1.).

Structure prediction of SalCis21 and SalCis20 identified a conserved HEXXH motif, demonstrating similarity to the C-terminal metalloprotease domain of the T6SS effector of *Pseudomonas aeruginosa* (PDB: 6H56) VgrG2b^37^, previously implicated in periplasmic toxicity in *E. coli*^37^. A conserved DUF4157 domain was also identified in SalCis20, previously found adjacent to eCIS-associated toxins targeting bacterial or fungal non-kin microbes^38,39^.

To test for toxicity, the putative effectors were cloned separately into a pET11a vector and expressed in a pLysS plasmid harbouring strain of *E. coli.* Periplasmic toxicity of SalCis21 in *E. coli*, was noted when expressed with an N-terminal *OmpA* gene fusion to direct SalCis21 to the periplasm^40^ (Extended Data Fig. 9a). SalCis21 was not toxic in the cytosol (Extended Data Fig. 9a, Wt-OmpA-SalCis21). Periplasmic toxicity was abolished by triple point mutation in the HEXXH motif (H99A, E100A, H103A), as well as by the E100A single substitution.

In contrast, SalCis20 was toxic upon periplasmic and cytosolic expression. HEXXH motif mutations (H150A, E151A, H154A), N- or C-terminal truncations, and a single E151A mutation failed to abolish toxicity (Extended Data Fig. 9b).

Co-expression of both effectors, expressed in tandem with and without the *OmpA* gene fusion did not abolish toxicity (Extended Data Fig. 9c), and no cognate immunity protein was identified. SalCis21 was less toxic compared to an established periplasmic toxin, Tse1 associated with the *P. aeruginosa* T6SS^41^ (Extended Data Fig. 9d). These results suggest that SalCIS encodes two toxic effectors, with SalCis20 utilizing an unknown mechanism of action, although we cannot rule out that the effect SalCis20 overexpression is non-specific and due to its non-native expression levels. Notably, the AlgoCIS effectors Cgo1 and Cgo2, both exhibited such toxicity, and similar HEXXH motifs^4^.

### Canonical eCIS-like secretion was not detected in SalCIS

We engineered a ΔSalCIS deletion strain of *S. enterica* subsp. *salamae*, and reintroduced the full operon on an inducible plasmid, to assess whether SalCIS functions as a canonical eCIS. Here, the SalCIS operon was expressed in tandem, as described previously. Upon induction, neither the sheath protein SalCis2, nor the two putative effectors, were detected in the extracellular fraction under tested conditions. They were only detected in the cellular fraction, probed via polyclonal antibodies using Western blot (Supplementary Fig. 4).

While regulated secretion cannot be excluded, absence of detectable proteins supports a non-canonical or potentially domesticated/defensive role, involving an alternative activation pathway. Taken together, the data suggest SalCIS diverges from known eCIS mode of action and may represent a distinct, conditionally regulated system.

### Cryo-ET reveals SalCIS interacts with the inner membrane

To investigate *in situ* localization, the operon was heterologously expressed in a minicell-producing strain of *S. typhimurium*. Here, the SalCIS operon was expressed in tandem; SalCis15, 20 and 21 were reversed and cloned to allow constitutive translation of all genes. The reconstructed tomograms revealed CIS localising close to the inner membrane of minicells (Fig. 5a). The number of CIS observed per bundle varied between 1-10, measuring between 600-700nm in length, and a uniform width of 18nm per CIS, congruent with the extended SalCIS diameter from SPA analysis. Notably, SalCIS expression in minicells led to arrested division at expected scission sites, assumedly hindered from budding by the SalCIS. The inner membrane appeared pinched when in the proximity of SalCIS (Fig. 5a). This localization is reminiscent of what has previously been described for the cryo-ET analysis of the *Amoebophilus asiaticus* T6SS^iv^.

**Figure 5.**
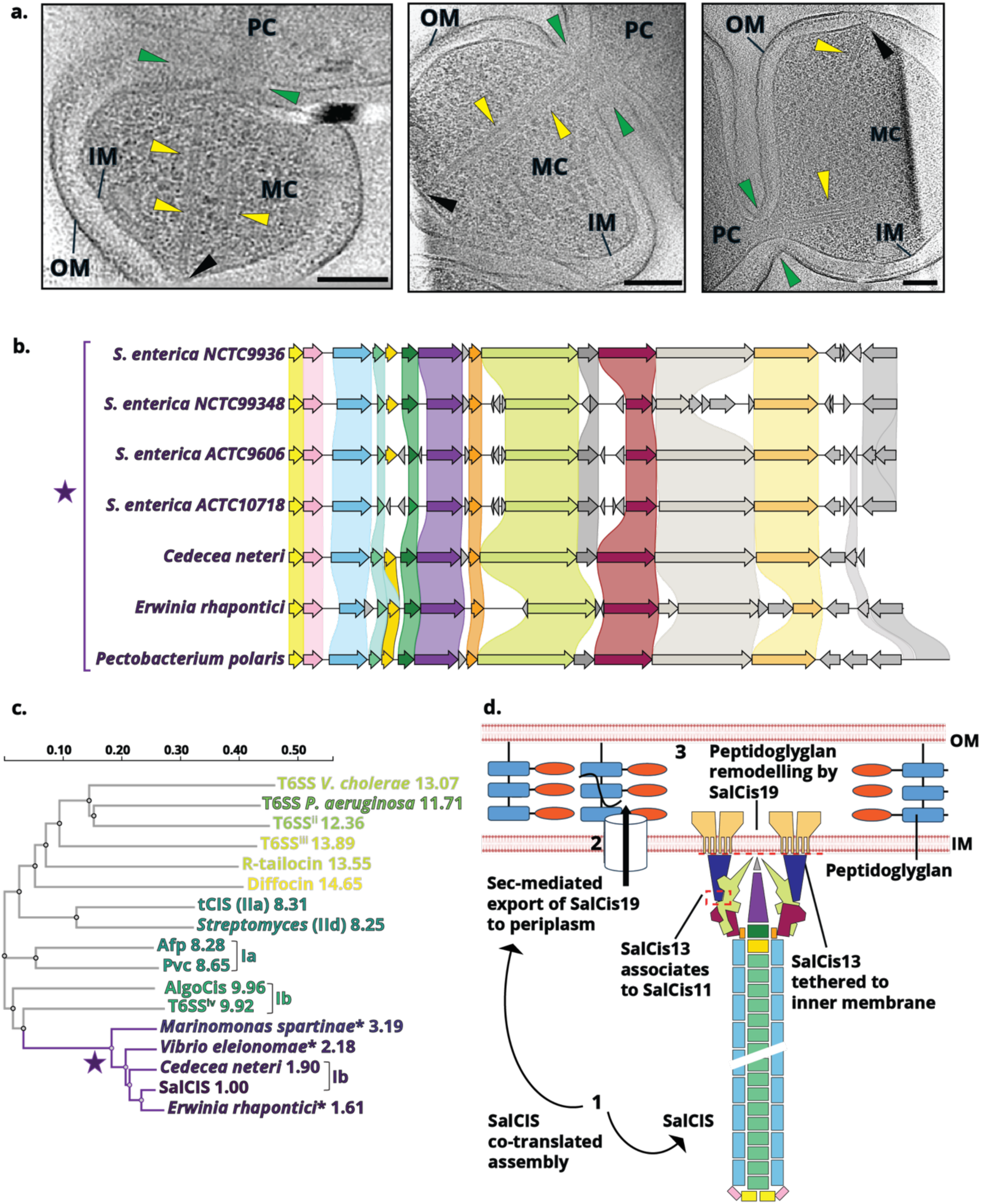
Cryo-ET localization and evolutionary context of SalCIS. **a:** Three individual, reconstructed cryo-electron tomogram slices, from three separate cells illustrating key features. Minicells (MC) are imaged emerging from the parent cell (PC). Green arrows indicate the expected sites of membrane scission during minicell formation. Yellow arrows highlight SalCIS bundles positioned within the cytoplasm. The inner (IM) and outer (OM) membranes of the Gram-negative *S. typhimurium* cell are labelled. The black arrow denotes potential interaction between SalCIS structures and the inner membrane. Black scale bar, 100 nm. **b:** Alignment of four SalCIS, and similar clusters in other bacteria employing Clinker^42^, forming a distinct cluster, genes are coloured according to SalCIS homology. **c:** Multiple sequence alignment (MSA) of CIS sheath proteins was performed using Clustal Omega^43^. The resulting guide tree was used to visualize evolutionary relationships. Branch lengths were extracted and normalized to the SalCis2 sheath protein (set as 1.0) to enable relative comparison of divergence. eCIS clade lineages are mentioned in brackets or grouped. Asterix indicates sheath proteins from unannotated eCIS operons identified in this study. **d:** Cartoon depiction of the postulated mode of membrane anchoring in SalCIS. Once all components of the operon are expressed, SalCis9 is exported to the periplasm via a Sec-mediated translocation pathway (1). The signal sequence cleaves as Sec-translocon recognition results in membrane embedded TMDs, followed by local remodelling of the peptidoglycan layer (2). SalCis19 tethers SalCis13 and the SalCIS baseplate to the membrane (3).

### SalCIS represents a unique cluster of CISs

Based on the unique membrane-associated structure of SalCis19, the predicted trimeric architecture of SalCis13, and the extended domain IV of SalCis11, we searched for related CIS operons containing homologues of these components. This analysis identified a distinct group of CISs that share conserved features (Fig. 5b). Phylogenetic comparison of their sheath proteins revealed a coherent cluster, suggesting evolutionary and functional relatedness (Fig. 5c) All operons in this cluster also encode a homologue of SalCis20, perhaps a core effector of this CIS subtype. Notably, SalCis21 is absent from any CIS cluster, including canonical eCIS and T6SS, and may be a lineage-specific toxin acquired horizontally by *S. enterica*. AlphaFold3 pull down of all proteins in the operon versus each other revealed predicted interactions supporting our models (Supplementary Fig. 5). Taken together, we hypothesize that SalCIS belongs to a structurally distinct subtype of CISs, potentially specialized for inner membrane association and peptidoglycan remodelling (Fig. 5d).

## Conclusion

In this study, we present the near-complete atomic structure of a previously uncharacterized extracellular contractile injection system (SalCIS) in *Salmonella enterica* subsp. *salamae.* Our data suggest that the SalCIS shares some core structural features with eCISs yet exhibits significant deviation from canonical eCIS and T6SS architecture. Saliently, unlike PVCs and Afps, SalCIS appears to operate as a membrane-associated complex, lacking detectable extracellular secretion of structural components or toxins.

Central to this divergence is the integration of a transmembrane protein, with a putative hydrolase domain, and the trimer adhesin-like SalCis13, within the core SalCIS operon. These proteins were structurally predicted to interface with the baseplate, suggesting a model in which SalCIS is tethered to the inner membrane rather than released extracellularly. Cryo-ET analyses further support this membrane association, though we could not definitively resolve anchoring orientation due to cell thickness.

We also identified two putative effectors within the operon. SalCis21 exhibits mild periplasmic toxicity, while SalCis20 was toxic in both periplasmic and cytosolic contexts in *E. coli*, suggesting a distinct but incompletely characterised mode of action. Their presence underscores the potential of SalCIS as a functionally active system, likely regulated in contexts beyond tested conditions.

Our findings pose SalCIS as an emerging class of CISs that appear domesticated, conditionally regulated for niche-specific interactions, such as competition or symbiosis. The large structural size of SalCIS and its membrane interaction offer promising avenues for future bioengineering efforts to deliver customized effectors in targeted drug delivery applications.

## Methods

### Bacterial strains

For this study, the CIS clusters of *Salmonella enterica* subsp. *salamae* serovar Greenside strains, NCTC9936, NCTC9948, *Salmonella enterica* subsp. *salamae* serotype Rowbarton, NCTC10718, NCTC9606 were procured from National Collection of Type Cultures (NCTC) and American Type Culture Collection (ACTC). For *in situ* studies, *Salmonella typhimurium* strain, ATCC 14028, mutated in *sseC* gene of the *Salmonella* pathogenicity island 2 (SPI-II), was used and further mutations were carried out in this strain background for cryo-ET and eCIS expression.

### Generation of *Salmonella* CIS plasmids and mutant strains

To obtain the atomic structure of the *Salmonella* CIS, the entire CIS cluster was cloned from *Salmonella enterica* subsp. *salamae* serovar Greenside strains, NCTC9936 and NCTC9948. The cluster was cloned as present on the bacterial genome, but with the last three genes reversed to induce expression in tandem. Genomic DNA was purified from overnight culture of *Salmonella* strains using GenElute kit (Thermofisher). Gene fragments were amplified using polymerase chain reaction (PCR) and cloned into linearized pBAD33 vector, using In-Fusion cloning (Takara Bio). Deletion clones of SalCis16b, SalCis13 and SalCis19 were obtained using PCR, followed by assembly with In-Fusion cloning. For spot assays, *S. enterica* CIS effector plasmids were cloned using gene synthesis and sub-cloning services from GenScript. For single particle analysis CIS plasmids were electroporated into electrocompetent BL21 Star^TM^ (DE3) cells (2.5 kV, 25 mF, 200 Ω). For cryo-electron tomography, CIS plasmids were transformed into electrocompetent *S. typhimurium* mutant strains.

Genomic mutants of *S. typhimurium* to induce mini cell expression, were generated using the λ Red system, wherein the *minD* was deleted to produce aberrant cell division resulting in mini cells. Genomic mutants of *S. enterica* subspecies *salamae* to delete SalCIS from the genome, were generated using the λ Red system too, wherein the system was deleted in parts.

Plasmid sequences were confirmed with whole plasmid nano-pore sequencing by Macrogen Europe.

### Purification of CIS for electron microscopy and LC-MS

E. coli BL21 StarTM (DE3) cells harbouring CIS plasmids were inoculated from a single colony on a plate into LB at 37°C for overnight preculture. Preculture was used to inoculate 2L of LB (1:100 ratio), cultivated at 37°C, until O.D.600 nm ≈1.0 and induced with L-arabinose for 24 h, induced cultures were then pelleted (5000g for 15 min). Pelleted cultures were resuspended in 100 mL cold Milli-Q water and pelleted again (5000*g* for 15 min). The pellet was then flash frozen, to break up cell walls, and subsequently thawed in a warm water bath, resuspended in 20 mL lysis buffer (50 mM Tris-HCl, 150 mM NaCl, 1% Triton X-100, 0.5x CellLytic Reagent (Sigma-Aldrich), 200 µg mL^-1^ lysozyme (in 10 mM Tris-HCl, pH 8.5), 50 µg mL^-1^ DNase I, 5 mM EDTA, 10 mM MgCl_2_, supplemented with protease inhibitor cocktail (Roche), pH 7.4) and incubated at 37 °C for 30 min. Cell debris was cleared (15,000*g* for 15 min). To clear remaining debris, the sample was spun for 30 min at 30,000*g*. To pellet CIS, the supernatant was spun for 1 h at 100,000*g*. Pelleted CIS were resuspended in 1 mL of resuspension (RS) buffer (50 mM Tris-HCl, 150 mM NaCl, supplemented with protease inhibitor, pH 7.4), by gentle agitation on a rotary roller at 4 °C for 1 h. The resuspended CIS were then loaded on a 10-40% iodixanol gradient, prepared in RS buffer and centrifuged at 100,000*g* for 24 hours. The gradient was fractionated, followed by SDS-PAGE and Western Blot analysis to confirm presence of CIS. Fractions containing CIS were pooled and dialysed for 4 days to remove iodixanol, at 4 °C. The dialysed CIS were then pelleted for 1 h at 100,000*g*, and used for negative stain TEM and cryo-EM. For LC-MS, the bidirectional operon was co-expressed, and purified in the same manner.

### Purification of CIS membrane component

For isolating the membrane bound SalCis19, 2L of *E. coli* BL21-Gold^TM^ cells harbouring plasmid with SalCis19 were cultivated and induced for 24 hours with IPTG at 16°C, 160rpm. Cultures were pelleted (5,000g, 20 min), flash frozen, and thawed, followed by resuspension in buffer (20 mM Tris-HCl, 150 mM NaCl, 10 mM MgCl_2_, 5 mM EDTA, 1 mM TCEP, and 10% glycerol, pH 7.5). Cells were lysed using a homogenizer (Avestin Emulsiflex C5). Cell debris was spun down (10,000g, 15 min), and any left-over debris was cleared by another round of centrifugation (30,000g, 30 min). Supernatant was supplemented with 10 mL of buffer and homogenized using a stirrer at 10°C for 2 hours. Cell membranes containing target proteins were pelleted (100,000g, 1hr). Membranes were then solubilized overnight at 10°C after adding Lauryl maltose neopentyl glycol (LMNG) to 1%. Strep-tag purification using a gravity drop set up was used to separate the strep-tagged transmembrane hydrolase and then eluted with elution buffer. Finally size exclusion chromatography was used to separate the target protein from the background. SalCis19 peak eluted at 300kDa (Superdex 200 GL column, GE Healthcare, equilibrated with buffer and 0.02% LMNG) and used for SDS-PAGE, Western blot analysis and cryo-EM grid preparation. To isolate SalCIS from endogenous expression along with the membrane component *S. enterica* subspecies *salamae* cells harbouring the wild-type SalCIS on a plasmid were cultivated and induced for 24 hours O.D._600_nm 1.0, with L-Arabinose at 18 °C, 60rpm. Cultures were pelleted (6000g, 20 min), flash frozen in liquid nitrogen, thawed and then resuspended in buffer (20 mM Tris-HCl, 150 mM NaCl, 10 mM MgCl_2_, 5 mM EDTA, 1 mM TCEP, and 10% glycerol, pH 7.5). This was followed by sonication (30% amplitude, 15s on/20s off pulses, 10 min on ice). Cell debris was pelleted at 13,000g, followed by 40 passes in a Dounce homogenizer. Membranes were solubilized in 1% DDM at 4°C. Membrane and CIS were pelleted through a 40% sucrose cushion at 150,000g for one hour. Resuspended pellet was cleaned by running through a 10-40% sucrose gradient, and fractionated, and then checked on TEM and Western blot.

### Negative-stain TEM

Concentrated CIS samples were spun at 10,000*g* to pellet aggregates. 4 μL of sample was applied to glow-discharged (15 mA, 45 s), continuous carbon Cu grids (200 mesh), for 60 s. The sample was washed twice with RS buffer and stained with 2% uranyl acetate for 60 s. Grids were inspected using a Thermo Fisher Scientific Morgagni transmission electron microscope operated at 80 kV.

### Single particle cryo-EM

Purified SalCIS particles were plunge-frozen using a Vitrobot^TM^ Mark IV (Thermo Fisher Scientific). 3.5 μL of sample was applied to glow discharged (5 mA, 10 s) R2/2 200 mesh (Quantifoil) Cu grids coated with an extra 2 nm layer of continuous carbon support film. Grids were blotted for 3s, blot force -8, and plunge frozen into liquid ethane, cooled by liquid nitrogen. 2.5 mg mL^-1^ of purified SalCis19 was applied to glow discharged (15 mA, 30 s) R0.6/1 200 mesh (Quantifoil) Au grids. Grids were blotted for 5s, blot force 15, and plunge frozen into liquid ethane, cooled by liquid nitrogen. For SalCIS particles data was collected from 2 grids using a Titan Krios G2 (Thermo Fisher Scientific), operating at 300 kV, equipped with an Selectris X energy filter (slit width 10 eV) and a Falcon 4i detector. Data was collected at a magnification of 165,000 (pixel size 0.728 Å) and target defocus range of 0.5 to 2.5 μm, and a total dose per acquisition of 41 e^−^ Å^-2^, fractionated in 41 frames, using EPU software. For SalCis19, data was collected from 1 grid as mentioned. To improve a slight orientation bias, data was also collected at -40 and +40 degrees stage tilt. Data were also collected from 1 grid using a Glacios Krios (Thermo Fisher Scientific), operating at 200 kV, and a Falcon 3 detector.

All data processing, classifications and reconstructions were carried out in cryoSPARC (Supplementary Fig. 6). For assembled SalCIS, two datasets of about 80,000 movies were patch motion corrected, followed by patch CTF estimation. Particles were picked using Blob Picker, and manually curated to remove bad particles, thick ice and foil edge images. Extracted particles were subjected to several rounds of 2D classification. Due to the heterogenous nature of the dataset, the same dataset was used to pick baseplates, caps and extended sheath assemblies. 2D classes chosen for each category were then used as templates to pick further particles, followed by one more round of 2D classification. After an initial reconstruction (C6 symmetry imposed), the particles were further classified using 3D classification, and bad particles rejected after manual inspection. For the cap, 700 particles were picked manually, and the initial reconstruction (C6 symmetry imposed) was used to generate 2D templates for template picking. Chosen volumes and their corresponding particles were used in homogenous refinement (C6 symmetry imposed), followed by re-extraction of aligned and per particle CTF corrected particles, reconstruction only, homogenous refinement, and a final round of non-uniform refinement (C6 and C3), to obtain maps with global resolutions of 3 Å (Supplementary Table 1). Local refinements using symmetry expanded particles were used to resolve densities at the edge of the baseplate (C6), cap (C6) and inside the central spike (C3), resulting in maps at a resolution of 2.86 Å and 2.97 Å, and 2.3 Å, respectively (Supplementary Fig. 6 and 7). For the contracted sheath, about 8060 movies were processed in the same manner, resulting in a map of 3.24 Å (supplementary Fig. 8). For the membrane protein SalCis19, from a dataset of about 42067 movies particles were picked and classified in the same way, with C2 symmetry imposed. Resulting in a map at 2.73 Å resolution (Supplementary Fig. 6 and 7).

### Atomic model building

All atomic model building was carried out by rigid-body fitting AlphaFold3-multimer predictions (Supplementary Fig. 9) into maps in ChimeraX^42^. The models were then fitted into the maps using the all-atom simulation in ISOLDE^43^ inside ChimeraX, and manually rebuilt. For the baseplate, two different maps were used to fit the C6 baseplate components, including SalCis11, SalCis12, SalCis2, and SalCis9. The maps used were non-sharpened C6 and C3 non-uniform refined reconstructions, as superimposition of various reconstructions revealed missing densities from different maps. Broken down densities towards the map edges required use of 2 separate local refinements using masks. Masks were segmented in ChimeraX using Segger segmentation tool^44^. For the spike puncture and inner tube initiation complex, SalCis10, SalCis8, SalCis6, SalCis7 and SalCis5, were similarly predicted, and fit and built onto the baseplate spike puncture complex map. Two cap map densities were used to fit and rebuild AlphaFold3 predicted models of SalCis16a, SalCis16b, and the terminal ends of sheath and inner tube complexes. For model building of the contracted sheath, the AlphaFold3 predicted model of SalCis2 and SalCis1 were fit into a helically refined map of the extended sheath, whereas only SalCis2 was built into the contracted sheath map and rebuilt using the same workflow. All models were then iteratively refined using real-space refinement in PHENIX^45^ (Supplementary Fig. 10). Portions of the predicted models could not be resolved in the map and were truncated in the final models (Supplementary Table 1.). MolProbity^46^ (PHENIX) was used to validate models (Supplementary Table 3). ChimeraX was used for graphical depiction of all models and maps.

### Cryo-ET sample preparation

*S.* Typhimurium cultures, heterologously expressing *S. enterica* subspecies *salamae* CIS were pelleted (2,000*g* for 20 min), and resuspended in cell resuspension (CRS) buffer (136mM NaCl, 2.5 mM KCl, 1.3mM MgCl2, 2mMCaCl2, 10mM HEPES, 10mM D-glucose, sterile filtered), and incubated at room temperature for 1 h. 2.5 μL of sample was applied to glow discharged (15 mA, 45 s) Quantifoil R2/2 Au 200 mesh (Jena Bioscience) grids, 0.5 μL of Protein-A 10 nm Gold Conjugate (Cytodiagnostics) was applied directly to the sample and gently mixed, followed by incubation for 1 min. The sample was then back blotted manually for 9 s, and plunged frozen into liquid ethane, cooled by liquid nitrogen, using a Vitrobot Mark IV (Thermo Fisher Scientific). Grids were stored in liquid nitrogen.

### Cryo-ET data collection

*S.* Typhimurium expressing CIS were loaded onto grids and cryo-ET data was collected, twice. The first collection was carried out at a magnification of 2.16 pixel size, using a Titan Krios G2 (Thermo Fisher Scientific), operating at 300 kV, FEG transmission equipped with Ceta-D camera and a Gatan K3 BioQuantum detector, with SerialEM software, at SciLife lab Stockholm, as a dose symmetric collection at 2° increments, and an angular range from -58° to +58°, with a defocus of -10 to -3 μm, and a total accumulated dose of 110 -120 e^−^ Å^-2^. The second data collection was carried out at Diamond Light Source, on a Titan Krios G3i (I) with BioQuantum LS imaging filters (slit width 20 eV) and K3 direct electron detectors (Gatan). Search maps at low magnification were used to identify target anchor points. Tomo5 software was used to automatically collect data in 2° or 3° increments, and an angular range from -60° to +60°, with a defocus of -9 to -7 μm. A total accumulated dose of 130 – 150 e^−^ Å^-2^, and a pixel size of 2.142 Å was used.

### Tomogram reconstruction and segmentation

Cryo-electron tomograms were reconstructed and segmented using the IMOD software suite^47^. Tilt-series alignment was performed using fiducial-based tracking with 10 nm protein-A gold markers, followed by CTF correction in Ctfplotter. Final reconstruction was carried out using weighted back-projection applying a SIRT-like filter (12). Reconstructed tomograms were visualized and manually annotated in 3dmod. Segmentation of SalCIS bundles was performed manually using 3dmod, following the tube segmentation strategy. Open contours were placed along the central axis of each SalCIS particle, followed by interpolation and surface generation to capture the sheath. Automated membrane segmentation was performed using MemBrain^48^. All segmented models were imported into ChimeraX for annotation, visualization, and figure generation. SalCIS bundles, membranes, and vesicles were coloured and labelled for clarity, and final images were exported for use in figures.

### Heterologous expression of SalCIS

For single particle cryo-EM, BL21 Star™(DE3) were used to heterologously express SalCIS. Strains were cultured in 2L of LB medium supplemented with appropriate antibiotics (100 *µ*g mL^−1^ ampicillin and 34 *µ*g mL^−1^ chloramphenicol) at 37 °C and constant agitation (180 r.p.m) until OD_600_ 0.8 was reached. This was followed by induction of SalCIS expression (0.2% w/v L-arabinose and/or 0.5mM isopropyl β-d-1-thiogalactopyranoside), continued incubation with agitation at 37 °C, until OD_600_ 1.2, and then slowly cooled down to 18 °C. SalCIS expression was induced for 24 or 48 h, under gentle agitation (70 r.p.m). For cryo-ET, *S. typhimurium* cells harbouring SalCIS plasmids were grown to OD_600_ 0.2 at 37 °C, under constant agitation (180 r.p.m), followed by induction of heterologous SalCIS expression with L-arabinose (0.2% w/v), until OD_600_ 0.8 was reached. Cells were then cooled to 18 °C, followed by induction for 24 h, under gentle agitation (70 r.p.m).). A BSL-1 strain of *S.* Typhimurium LT2 was used for cryo-ET, as our imaging and cryo-EM sample preparation facility is not biosafety level 2 designated.

### Secretion assays and Western blotting

*Salmonella enterica* subspecies *salamae* strain with a deletion of SalCIS (ΔSalCIS) was transformed with SalCIS genes. ORFs 1-15 (SalCis16a to SalCis19) were cloned on pBAD33 (L-Arabinose inducible), while ORFs 16-18 (SalCis15-SalCis21) were cloned onto pWSK29 (low copy number, IPTG-inducible). Strain and controls were cultured in LB medium with appropriate antibiotics at 37 °C, 180 rpm. Overnight cultures were diluted 1:1000 in 5 mL LB and grown for 1.5 h, followed by induction of gene expression with 2% L-arabinose and 0.2 mM IPTG for 2 h. Cultures were then centrifuged to separate cell pellets from supernatants. Secreted proteins in the supernatant were precipitated using trichloroacetic acid (TCA) and resuspended alongside cellular pellets in 2 x SDS sample buffer, normalized to 20 O.D. units per μL. Proteins were resolved by SDS-PAGE and transferred to nitrocellulose membranes for Western blot analysis. Detection was performed using anti-SalCis2, SalCis1, SalCis20 and SalCis21 polyclonal antibodies (0.2 mg mL^-1^ each). Cytosolic contamination was assessed using anti-DnaK antibody.

### LC-MS data

Purified SalCIS (100 µg) samples were diluted in 200 µL 50 mM TRIS pH 8.5 at room temperature and split into two equal aliquots. Each aliquot was then processed by one of two digestion protocols: in-solution digestion or Protein Aggregation Capture (PAC). In each case, samples were digested in parallel with either sequencing-grade trypsin (Promega), or endoproteinase Lys-C (Wako). For in-solution digestion, 0.25 µg of proteolytic enzymes were added and samples incubated overnight at 25°C. For PAC, proteins were precipitated by adding 10 volumes of acetonitrile, immobilized on magnetic microparticles. The microparticles were fixed using a magnet stand, washed twice with 100% acetonitrile and once with 70% ethanol, after which microparticles were air-dried for 15 min, removed from the magnet stand, and resuspended in 50 mL 50 mM TRIS pH 8.5. For subsequent digestion, 0.25 µg of proteolytic enzymes were added, and samples were incubated overnight at 30 °C shaking at 1,250 RPM, after which magnetic microparticles were fixed using a magnet stand and the digests transferred to new tubes. For both digestion strategies, samples were reduced and alkylated by concomitant addition of tris(2-carboxyethyl)phosphine and chloroacetamide to final concentrations of 10 mM, followed by incubation at 30 °C for 30 min. All samples were clarified through 0.45 µm spin filters, and peptides were purified via low-pH C18 StageTip procedure. To this end, C18 StageTips were prepared in-house, by layering four plugs of C18 material (Sigma-Aldrich, Empore SPE Disks, C18, 47 mm) per StageTip. Activation of StageTips was performed with 100 μL 100% methanol, followed by equilibration using 100 μL 80% acetonitrile (ACN) in 0.1% formic acid, and two washes with 100 μL 0.1% formic acid. Samples were acidified to pH < 3 by addition of trifluoroacetic acid to a final concentration of 1% (vol/vol), and loaded on the StageTips. Subsequently, StageTips were washed twice using 100 μL 0.1% formic acid, after which peptides were eluted using 80 µL 25% ACN in 0.1% formic acid. The samples were dried to completion using a SpeedVac at 60 °C. Dried peptides were dissolved in 20 μL 0.1% formic acid (FA) and stored at −20 °C until analysis using mass spectrometry.

The four sample digests were analysed on a Vanquish™ Neo UHPLC system (Thermo) coupled to an Orbitrap™ Astral™ mass spectrometer (Thermo), as three technical replicates (1 µL, 2uL, and 3 µL injections). Samples were analysed on 15 cm long analytical columns, with an internal diameter of 75 μm, and packed in-house using ReproSil-Pur 120 C18-AQ 1.9 µm beads (Dr. Maisch). Elution of peptides from the column was achieved using a gradient ranging from buffer A (0.1% formic acid) to buffer B (80% acetonitrile in 0.1% formic acid), at a flow rate of 250 nL/min. Gradient length was 30 min per sample, including ramp-up and wash-out, with an analytical gradient of 20 min ranging in buffer B from 5-38%. The column was heated to 45 °C using a column oven, and ionization was achieved using a NanoSpray Flex™ NG ion source (Thermo). Spray voltage set at 2 kV, ion transfer tube temperature to 275°C, and RF lens to 50%. All full precursor (MS1) scans were acquired using the Orbitrap™ mass analyser, while all tandem fragment (MS2) scans acquired in parallel using the Astral™ mass analyser. Full scan range was set to 300-1,300 m/z, MS1 resolution to 120,000, MS1 AGC target to “250” (2,500,000 charges), and MS1 maximum injection time to 150 ms. Precursors were analysed in data-dependent acquisition (DDA) mode, with charges 2-6 selected for fragmentation using an isolation width of 1.3 m/z and fragmented using higher-energy collision disassociation (HCD) with normalized collision energy of 25. Monoisotopic Precursor Selection (MIPS) was enabled in “Peptide” mode. Repeated sequencing of precursors was minimized by setting expected peak width to 10 s, and dynamic exclusion duration to 7.5 s, with an exclusion mass tolerance of 10 ppm and exclusion of isotopes. MS2 scans were acquired using the Astral mass analyser. MS2 fragment scan range was set to 100-1,500 m/z, MS2 AGC target to “100” (10,000 charges), MS2 intensity threshold to 100,000 charges per second, and MS2 maximum injection time to 5 ms.

### LC-MS Data analysis

MS RAW data were analysed using MaxQuant software (v.2.4.3.0). Default MaxQuant settings were used, with exceptions specified below. For generation of theoretical spectral libraries, in-silico digestion was performed on a database containing 19 expected protein sequences from SalCIS. In-silico digestion was performed with trypsin, in semi-specific digestion mode, with minimum peptide length set to 6 and maximum peptide length set to 55. Allowed variable modifications were oxidation of methionine (default), protein N-terminal acetylation (default), deamidation of glutamine and asparagine, and peptide N-terminal conversion of glutamine to pyroglutamate, with maximum variable modifications per peptide set to 3. Modified peptides were filtered for a minimum score of 100 and a minimum delta score of 50. Second peptide search was disabled. Label-free quantification (LFQ) was enabled, with “Fast LFQ” disabled. Stringent MaxQuant 1% FDR data filtering at the PSM- and protein-levels was applied (default).

### SDS-PAGE and Western blotting

Samples were diluted and denatured for 5 min at 95°C in Laemmli buffer (Thermo Fisher Scientific), and then loaded on to 4-12% gradient Bis-Tris precast protein gels (Thermo Fisher Scientific). Gel electrophoresis was carried out at 200V in 1x MES running buffer for 35 min. Gels were stained with Coomassie blue. Gels for western blotting were run as described above, followed by transfer to nitrocellulose membranes (iBlot 2, Transfer Stacks, Invitrogen) using the iBlot system (iBlot 2 Dry Blotting System; Invitrogen) for 6 min. The membranes were then transferred to a sequential lateral flow (SLF), blocked with iBind solution kit (Invitrogen), incubated with primary and secondary antibodies (1/1000 dilution) from 3 hours to overnight. The blots were then visualized using TMB-D blotting solution (Kementec).

### Toxicity assays

The two putative effectors in the *S. enterica* CIS were cloned into pET11a (ampicillin resistance), codon optimized for *E. coli*. The *OmpA* periplasmic translocation sequence was fused to the N-terminus of the effectors, individually and in tandem. As controls, non-fusion effectors were expressed, in addition to, *Pseudomonas aeruginosa* Tse1 and sfGFP *OmpA* fusions. For mutational analysis, HEXXH motif and point mutants (e100a and e151a) were generated using PCR. Plasmids were then transformed to chemically competent Bl21 (DE3) pLysS cells using heat shock (42°C for 45 s). Overnight cultures were started from single colonies in LB medium supplemented with ampicillin, chloramphenicol and 1% D-glucose. After 20 h, overnight cultures were normalized to OD_600_ 0.8, pelleted at 6000*g* and resuspended in fresh LB medium. Serial dilutions of cultures were then spotted onto plates under induction (0.5mM IPTG) and suppression (1% D-glucose) conditions and incubated at 37°C for 24 h. Plates were imaged using a plate scanner.

### Bioinformatic analysis and structure prediction

The cluster was identified employing the eCIS-screen script ^1^, whereas the start and end of the putative cluster was confirmed by presence of natural Shine-Dalgarno sequences (AGGAGG and GAGG). HHPred was used inside the MPI Toolkit, to search for protein sequence based structural homology of all proteins in the cluster ^49^. AlphaFold3-Multimer was used to predict structures of all proteins in the CIS cluster, for all four *S. enterica* strains^50^. The predicted structures were then submitted to the DALI server (Ekhidna 2 biocenter) to survey for structurally homologous proteins^51^. Transmembrane helixes were predicted using the DeepTMHMM 1.0 server^52^. SignalP 6.0 was employed to predict putative signal sequences in the SalCIS cluster^26^. Clinker was used to conduct operon alignments, and visualize gene cluster comparisons^53^. Electrostatic charge distribution computed using the Adaptive Poisson– Boltzmann Solver (APBS) at pH 7.4, with default dielectric values, molecular surfaces are coloured according to potential, from –5 kT/e (red) to +5 kT/e (blue)^54^. The all-vs-all AlphaFold3 predictions and code to plot Supplementary Figure 5 and Extended Data Figure 8b were carried out using custom scripts available at https://github.com/FJ0M/SalCIS_AF3_allvall. For the all-vs-all analysis, 5 models were created at 5 random seeds, creating 25 models for analysis. The value shown in the heatmap is the mean average of the scores calculated for the 25 models.

## Data availability

The cryo-EM density maps and corresponding atomic models have been deposited in the PDB and EMDB, respectively. The accession numbers are listed as follows: Baseplate module in extended state, reconstructed in C3 symmetry (PDB: 9RCE and EMD-53919); Cap module in extended state reconstructed in C6 symmetry (PDB: 9R9N and EMD-53859); Sheath and inner tube module in extended state (PDB: 9R9H and EMD-53857); Sheath in contracted state (PDB: 9R9I and EMD-53858); The membrane protein, SalCis19 as a dimer in LMNG (PDB: 9R9A and EMD-53855). The mass spectrometry data of purified SalCIS have been deposited to the ProteomeXchange Consortium, with the dataset identifier PXD063506.

## Acknowledgements

The Novo Nordisk Foundation Centre for Protein Research is supported financially by the Novo Nordisk Foundation (NNF14CC0001). This work was supported by a grant from the Independent Research Fund Denmark (1030-00289B). We thank the Protein Production Platform at CPR, (especially lab technician Michael Williamson), for their support in cloning large plasmids; staff at the Core Facility for Integrated Microscopy (especially TEM specialist Michael James Johnson) for their support; Diamond Light Source for time on Krios I, under proposal bi38819, at Harwell United Kingdom, and especially the late Dr. Karen Davies for her support in cryo-ET data collection; Dr. Simon Erlendsson for consultation on cryo-ET sample preparation; Professor Lone Brønsted at Copenhagen University for BSL-2 laboratory support (PHAGE group); Members of the Taylor group, including Nicole Rutbeek for consultation on membrane protein purification; Members of the Kummer and Montoya groups at the Novo-Nordisk Centre for Protein Research, and Dr. Magnus Bloch at Birkbeck University of London, for discussions. N.M.I.T. is a member of ISBUC. M.E. acknowledges funding by the Max Planck Society as Max Planck Fellow.

## Contributions

N.M.I.T and R.N.E conceived the project. C.S.K performed bioinformatic analysis. R.N.E. performed bioinformatic analysis, designed and cloned primary plasmid constructs, purified and prepared samples. R.N.E performed cryo-EM sample preparation. T.H.P and N.H.S collected cryo-EM data. R.N.E processed cryo-EM data, reconstructed cryo-EM maps, built, and refined the structures. R.N.E performed cryo-ET sample preparation and collected cryo-ET data with E.M.S.R. R.N.E processed, reconstructed, analysed and annotated tomograms. R.N.E designed and performed toxicity assays. F.J.O.M performed AlphaFold3 predictions, analysis, and bioinformatic analysis. K.F cloned all plasmids required for and designed, and performed genetic manipulations, genome editing, secretion assays and subsequent analysis in *S.* Typhimurium and *S. enterica* subspecies *salamae*. I.A.H designed and performed LC-MS and subsequent analysis. M.S prepared and uploaded structures to EMDB and PDB. R.N.E wrote the manuscript; all authors commented on the manuscript.

## Corresponding author

Correspondence to Nicholas M. I. Taylor

## References

1. Chen, L. et al. Genome-wide Identification and Characterization of a Superfamily of Bacterial Extracellular Contractile Injection Systems. Cell Rep. 29, 511–521.e2 (2019).

2. Lin, L. The expanding universe of contractile injection systems in bacteria. Curr. Opin. Microbiol. 79, 102465 (2024).

3. Weiss, G. L. et al. Structure of a thylakoid-anchored contractile injection system in multicellular cyanobacteria. Nat. Microbiol. 7, 386–396 (2022).

4. Xu, J. et al. Identification and structure of an extracellular contractile injection system from the marine bacterium Algoriphagus machipongonensis. Nat. Microbiol. 7, 397–410 (2022).

5. Jiang, F. et al. Cryo-EM Structure and Assembly of an Extracellular Contractile Injection System. Cell 177, 370–383.e15 (2019).

6. Taylor, N. M. I., Van Raaij, M. J. & Leiman, P. G. Contractile injection systems of bacteriophages and related systems. Mol. Microbiol. 108, 6–15 (2018).

7. Kikuchi, Y. & King, J. Genetic control of bacteriophage T4 baseplate morphogenesis. J. Mol. Biol. 99, 645–672 (1975).

8. Taylor, N. M. I. et al. Structure of the T4 baseplate and its function in triggering sheath contraction. Nature 533, 346–352 (2016).

9. Heymann, J. B. et al. Three-dimensional structure of the toxin-delivery particle antifeeding prophage of Serratia entomophila. J. Biol. Chem. 288, 25276–25284 (2013).

10. Evans, R. & Waterfield, N. R. The Pvc15 D2-Pnf SP Interaction Mediates AAA ATPase Activity, Payload Stability, and Translocation into the PVC Tube Lumen. Preprint at 10.1101/2023.08.15.553202 (2023).

11. Heiman, C. M., Vacheron, J. & Keel, C. Evolutionary and ecological role of extracellular contractile injection systems: from threat to weapon. Front. Microbiol. 14, 1264877 (2023).

12. Kreitz, J. et al. Programmable protein delivery with a bacterial contractile injection system. Nature 616, 357–364 (2023).

13. Vlisidou, I. et al. The Photorhabdus asymbiotica virulence cassettes deliver protein effectors directly into target eukaryotic cells. eLife 8, e46259 (2019).

14. Steiner-Rebrova, E. M. et al. N-terminal toxin signal peptides efficiently load therapeutics into a natural nano-injection system. Preprint at 10.1101/2023.11.13.566157 (2023).

15. Billah, M. M. & Rahman, M. S. Salmonella in the environment: A review on ecology, antimicrobial resistance, seafood contaminations, and human health implications. J. Hazard. Mater. Adv. 13, 100407 (2024).

16. Ofori, L. A. et al. Salmonella enterica in farm environments in the Ashanti Region of Ghana. BMC Microbiol. 23, 370 (2023).

17. Parsons, S. K., Bull, C. M. & Gordon, D. M. Spatial Variation and Survival of Salmonella enterica Subspecies in a Population of Australian Sleepy Lizards (Tiliqua rugosa). Appl. Environ. Microbiol. 81, 5804–5811 (2015).

18. Yan, M., et al. *Salmonella enterica* subsp. II serovar 4,5,12:a:-may cause gastroenteritis infections in humans. Gut Microbes 14, 2089007 (2022).

19. Bhardwaj, P., Mitra, A. K. & Hurst, M. R. H. Investigating the Process of Sheath Maturation in Antifeeding Prophage: a Phage Tail-Like Protein Translocation Structure. J. Bacteriol. 203, e0010421 (2021).

20. Desfosses, A. et al. Atomic structures of an entire contractile injection system in both the extended and contracted states. Nat. Microbiol. 4, 1885–1894 (2019).

21. Böck, D. et al. In situ architecture, function, and evolution of a contractile injection system. Science 357, 713–717 (2017).

22. Casu, B., Sallmen, J. W., Schlimpert, S. & Pilhofer, M. Cytoplasmic contractile injection systems mediate cell death in Streptomyces. Nat. Microbiol. 8, 711–726 (2023).

23. Shneider, M. M. et al. PAAR-repeat proteins sharpen and diversify the type VI secretion system spike. Nature 500, 350–353 (2013).

24. Yap, M. L. & Rossmann, M. G. Structure and Function of Bacteriophage T4. Future Microbiol. 9, 1319–1327 (2014).

25. Beveridge, T. J. Structures of Gram-Negative Cell Walls and Their Derived Membrane Vesicles. J. Bacteriol. 181, 4725–4733 (1999).

26. Nielsen, H., Teufel, F., Brunak, S. & Von Heijne, G. SignalP: The Evolution of a Web Server. in Protein Bioinformatics (ed. Lisacek, F.) vol. 2836 331–367 (Springer US, New York, NY, 2024).

27. Pradel, N., Delmas, J., Wu, L. F., Santini, C. L. & Bonnet, R. Sec- and Tat-Dependent Translocation of β-Lactamases across the *Escherichia coli* Inner Membrane. Antimicrob. Agents Chemother. 53, 242–248 (2009).

28. Üstok, F. I., Chirgadze, D. Y. & Christie, G. Structural and functional analysis of SleL, a peptidoglycan lysin involved in germination of *B acillus* spores: Crystal Structures for the Bacillus Spore Lytic Enzyme SleL. Proteins Struct. Funct. Bioinforma. 83, 1787–1799 (2015).

29. Tidhar, A., Rushing, M. D., Kim, B. & Slauch, J. M. Periplasmic superoxide dismutase SODCI of *S almonella* binds peptidoglycan to remain tethered within the periplasm. Mol. Microbiol. 97, 832–843 (2015).

30. Byun, B. et al. Mechanism of the Escherichia coli MltE lytic transglycosylase, the cell-wall-penetrating enzyme for Type VI secretion system assembly. Sci. Rep. 8, 4110 (2018).

31. Ashkenazy, H. et al. ConSurf 2016: an improved methodology to estimate and visualize evolutionary conservation in macromolecules. Nucleic Acids Res. 44, W344–W350 (2016).

32. English, G. et al. New secreted toxins and immunity proteins encoded within the T ype VI secretion system gene cluster of *S erratia marcescens*. Mol. Microbiol. 86, 921–936 (2012).

33. Sibinelli-Sousa, S. et al. A Family of T6SS Antibacterial Effectors Related to l,d-Transpeptidases Targets the Peptidoglycan. Cell Rep. 31, 107813 (2020).

34. Monchois, V., Abergel, C., Sturgis, J., Jeudy, S. & Claverie, J.-M. Escherichia coli ykfE ORFan Gene Encodes a Potent Inhibitor of C-type Lysozyme. J. Biol. Chem. 276, 18437–18441 (2001).

35. Rapisarda, C., et al. *In situ* and high-resolution cryo-EM structure of a bacterial type VI secretion system membrane complex. EMBO J. 38, e100886 (2019).

36. Caulton, S. G. et al. Bdellovibrio bacteriovorus uses chimeric fibre proteins to recognize and invade a broad range of bacterial hosts. Nat. Microbiol. 9, 214–227 (2024).

37. Wood, T. E. et al. The Pseudomonas aeruginosa T6SS Delivers a Periplasmic Toxin that Disrupts Bacterial Cell Morphology. Cell Rep. 29, 187–201.e7 (2019).

38. Geller, A. M. et al. The extracellular contractile injection system is enriched in environmental microbes and associates with numerous toxins. Nat. Commun. 12, 3743 (2021).

39. Danov, A. et al. Identification of novel toxins associated with the extracellular contractile injection system using machine learning. Mol. Syst. Biol. 20, 859–879 (2024).

40. Karyolaimos, A. & de Gier, J.-W. Strategies to Enhance Periplasmic Recombinant Protein Production Yields in Escherichia coli. Front. Bioeng. Biotechnol. 9, 797334 (2021).

41. Russell, A. B. et al. Type VI secretion delivers bacteriolytic effectors to target cells. Nature 475, 343–347 (2011).

42. Pettersen, E. F. et al. UCSF Chimera—A visualization system for exploratory research and analysis. J. Comput. Chem. 25, 1605–1612 (2004).

43. Croll, T. I. *ISOLDE* : a physically realistic environment for model building into low-resolution electron-density maps. Acta Crystallogr. Sect. Struct. Biol. 74, 519–530 (2018).

44. Pintilie, G. D., Zhang, J., Goddard, T. D., Chiu, W. & Gossard, D. C. Quantitative analysis of cryo-EM density map segmentation by watershed and scale-space filtering, and fitting of structures by alignment to regions. J. Struct. Biol. 170, 427–438 (2010).

45. Afonine, P. V. et al. Towards automated crystallographic structure refinement with *phenix.refine*. Acta Crystallogr. D Biol. Crystallogr. 68, 352–367 (2012).

46. Williams, C. J. et al. MolProbity: More and better reference data for improved all-atom structure validation. Protein Sci. 27, 293–315 (2018).

47. Kremer, J. R., Mastronarde, D. N. & McIntosh, J. R. Computer Visualization of Three-Dimensional Image Data Using IMOD. J. Struct. Biol. 116, 71–76 (1996).

48. Lamm, L. et al. MemBrain v2: an end-to-end tool for the analysis of membranes in cryo-electron tomography. Preprint at 10.1101/2024.01.05.574336 (2024).

49. Soding, J., Biegert, A. & Lupas, A. N. The HHpred interactive server for protein homology detection and structure prediction. Nucleic Acids Res. 33, W244–W248 (2005).

50. Varadi, M. et al. AlphaFold Protein Structure Database: massively expanding the structural coverage of protein-sequence space with high-accuracy models. Nucleic Acids Res. 50, D439–D444 (2022).

51. Holm, L. Dali server: structural unification of protein families. Nucleic Acids Res. 50, W210–W215 (2022).

52. Hallgren, J. et al. DeepTMHMM predicts alpha and beta transmembrane proteins using deep neural networks. Preprint at 10.1101/2022.04.08.487609 (2022).

53. Gilchrist, C. L. M. & Chooi, Y.-H. clinker & clustermap.js: automatic generation of gene cluster comparison figures. Bioinformatics 37, 2473–2475 (2021).

54. Jurrus, E. et al. Improvements to the APBS biomolecular solvation software suite. Protein Sci. 27, 112–128 (2018).

