## Extended Data for "Structure of a contractile injection system in *Salmonella enterica* subsp. *salamae*"

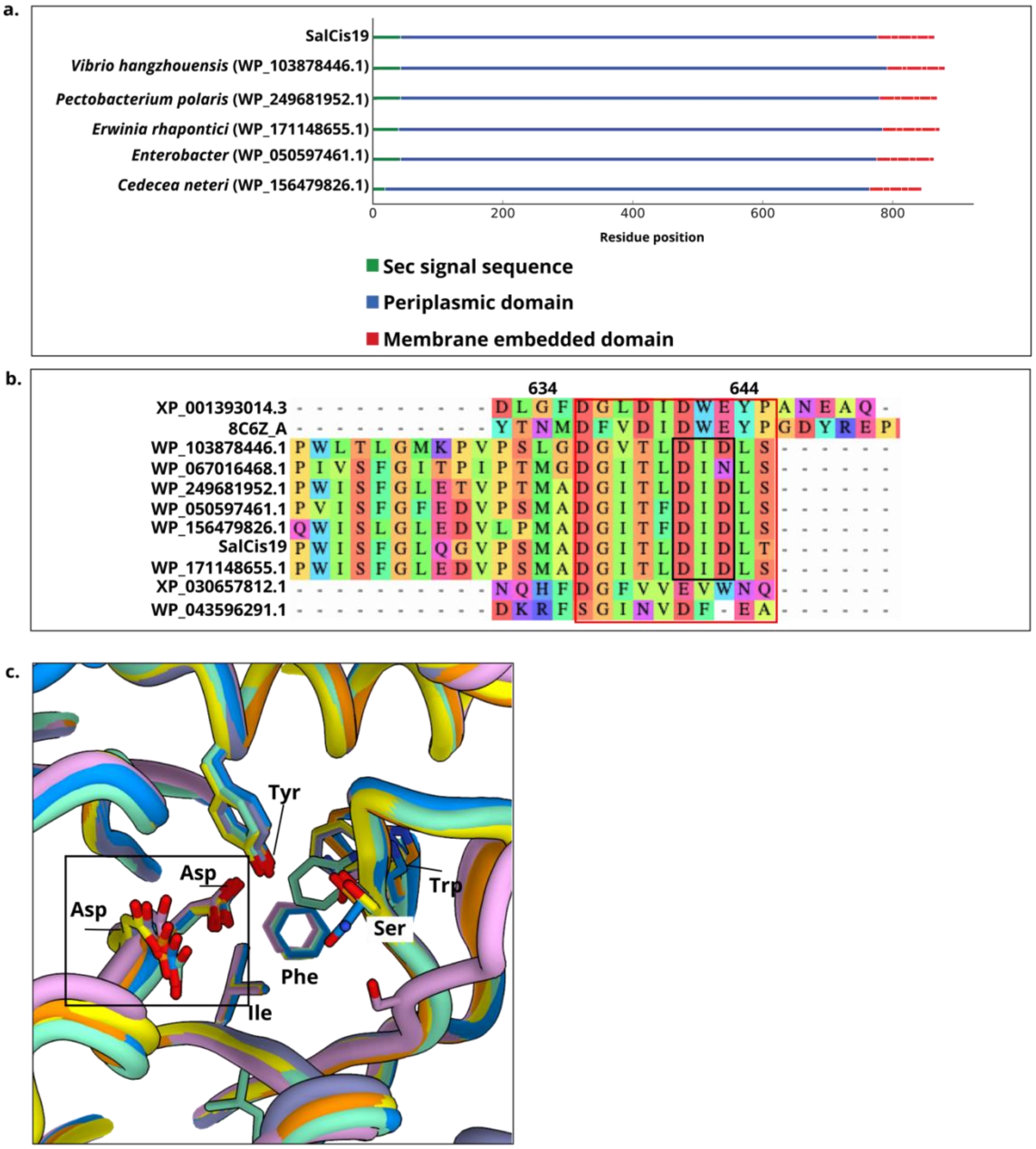


**Extended Data Figure 1: Comparison of SalCis19 homologues highlights similar topologies and a conserved enzymatic hydrolytic core**

**a:** Graphical depiction of topologies of selected SalCis19 homologues, homologues were detected using the Consurf server, and then the topologies were predicted using DeepTMHMM 1.0 server[52]. Species or taxonomy level names are followed by NCBI protein IDs in brackets. **b:** Multiple sequence alignment of SalCis19 homologues and related hydrolase enzymes. Red box indicates highly conserved enzymatic hydrolytic core across species, including bacteria (*Aspergillus niger*; XP_001393014.3, *Clostridium perfringens*; 8C6Z_A, *Chromobacterium violaceum*; WP_043596291.1) and mammals (*Nomascus leucogenys*; XP_030657812.1) **c:**

Structural alignment of AlphaFold3 structure predictions of SalCis19 homologues in a., with salient residues highlighted, black box indicates DXD motif conserved according to MSAs

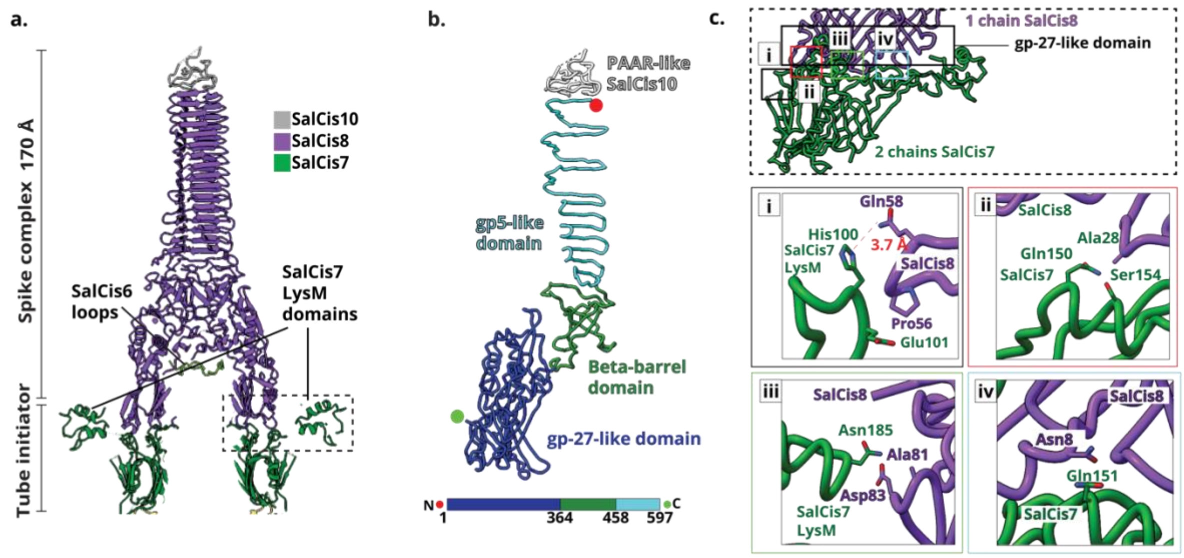


**Extended Data Figure 2: Extended details of the spike puncturing complex**

**a:** Cartoon depiction of SalCIS inner spike and tube initiator complex SalCis8. **b:** Domain arrangement of the SalCis8 spike, highlighting conserved domains. **c:** Inset details of black dashed box from panel a. Cartoons shows featured interacting residues. (i) His100 and Glu101 from SalCis7 contact Gln 58 in SalCis8. (ii) Gln150 and Ser154 from SalCis7 contact Ala28 from SalCis8. (iii) Asn185 from SalCis7's LysM domain contact SalCis8 Asp83 and Ala81. (iv) Gln151 from SalCis7 contact Asn8 from SalCis8.

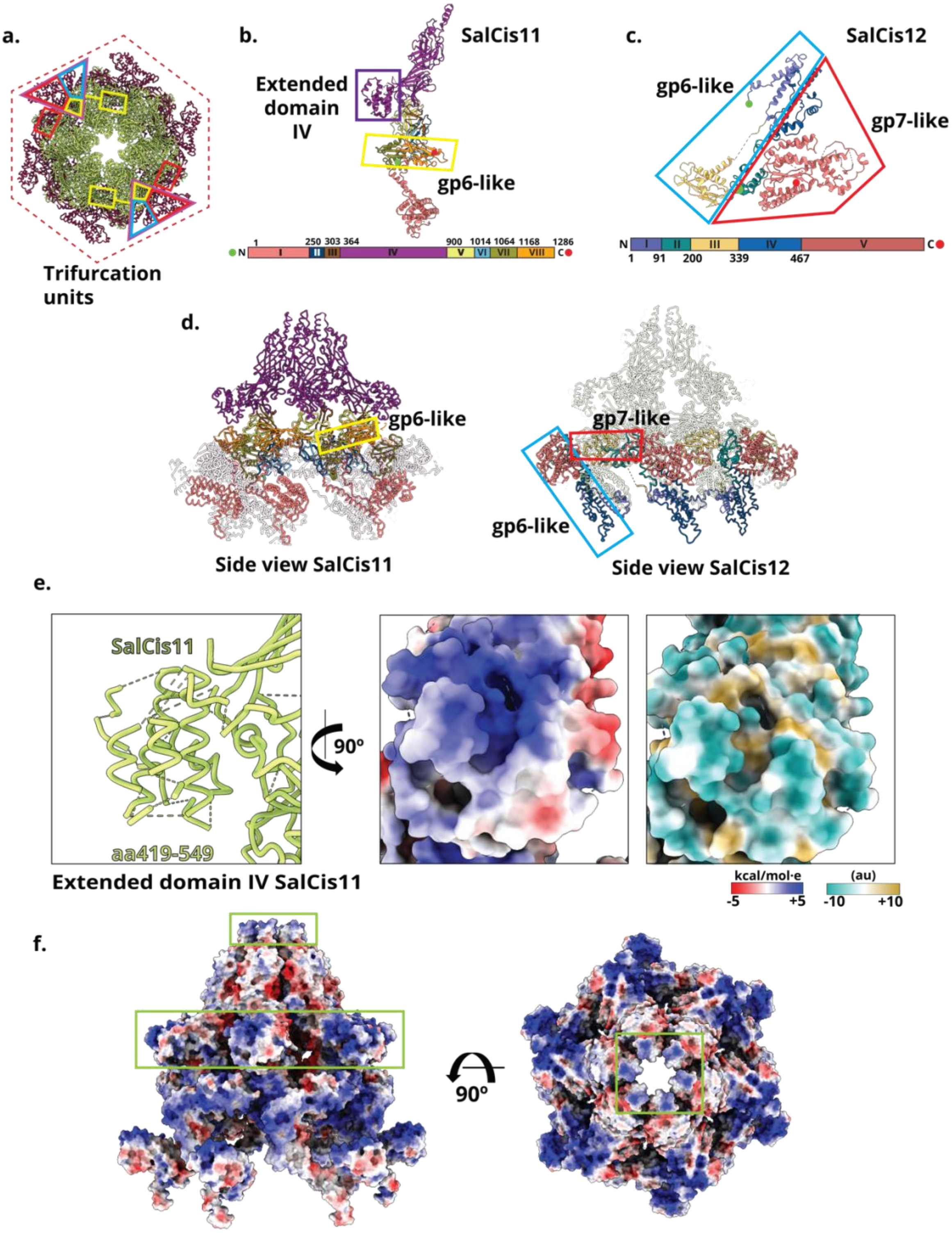


**Extended Data Figure 3: Extended details of the SalCIS cage and components**

**a:** Two conserved trifurcation units on the SalCIS baseplate are highlighted, as viewed from top of the cage with the compact hexagonal wedge highlighted. Red dashed hexagon highlights compact shape. **b:** Domain arrangement of SalCis11. Purple box indicates extension of domain IV unique to SalCIS. Yellow box indicates location of gp6-like domain. **c:** Domain arrangement of SalCis12, red outline indicates gp7-like features, blue box indicates gp6-like features. **d:** Right panel shows side view of the baseplate cage, with SalCis11 domain arrangement highlighted. Blue box indicates location of gp6-like features in the baseplate cage formed by SalCis11. Left panel shows side view of the baseplate cage, with SalCis12 domain arrangement highlighted. Green box indicates location of gp7-like features in the baseplate cage formed by SalCis12, black box indicates gp6-like domains. **e:** Details of SalCis11 and SalCis12. (i) Left panel shows cartoon of extended domain IV (ii) Middle panel shows coulombic electrostatic charge distribution, with a distinct positive charge distribution over the top surface of the extension (iii) Right panel shows hydrophobicity depiction of the same, highlighting hydrophilic residues over the outside of the extension, calculated in arbitrary units, Kyte-Doolittle scale. **f:** Electrostatic surface potential over SalCIS cage formation, showing six copies of SalCis11. Green box highlights the positively charge ring at the tip of the cage. Electrostatic charge distribution computed using the Adaptive Poisson–Boltzmann Solver (APBS) at pH 7.4, with default dielectric values, molecular surfaces are coloured according to potential, from –5 kT/e (red) to +5 kT/e (blue)[53].

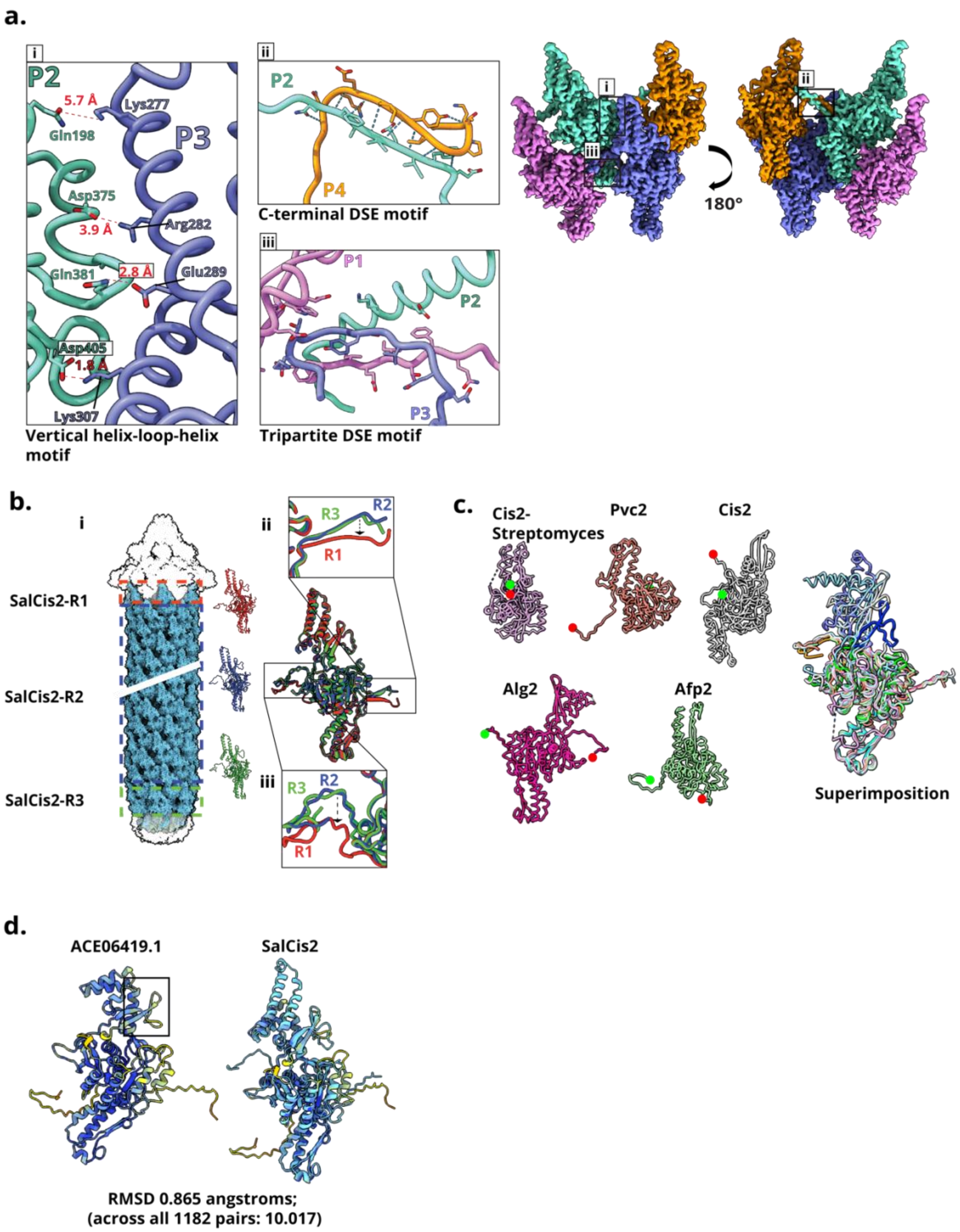


**Extended Data Figure 4. Extended details of outer sheath assembly and comparisons to other systems.**

**a:** Four protomers are highlighted and color-coded; coloured boxes indicate zoom-ins of key inter-protomer contacts, with corresponding cryo-EM density shown. (i) Shows distinct helix-loop-helix orientation of SalCis2, salient contacts and atomic distances are marked. (ii) Shows conserved C-terminus DSE motif. (iii) Shows a tripartite DSE motif, previously noted in tCIS.

**b:** Distribution of SalCis2 throughout the SalCIS sheath vertex, denoting three distinct conformational states. **c:** Structural comparison of SalCis2 with sheath proteins from other CISs, including superimposition to highlight variation between systems, and the X-loop of SalCis2. **d:** AlphaFold3 multimer predictions of a protein homologue of SalCis2 in *Amoebophilus asiaticus*. Black box in a. indicates the presence of the X-loop in the predicted model of the *A. asiaticus* sheath protein

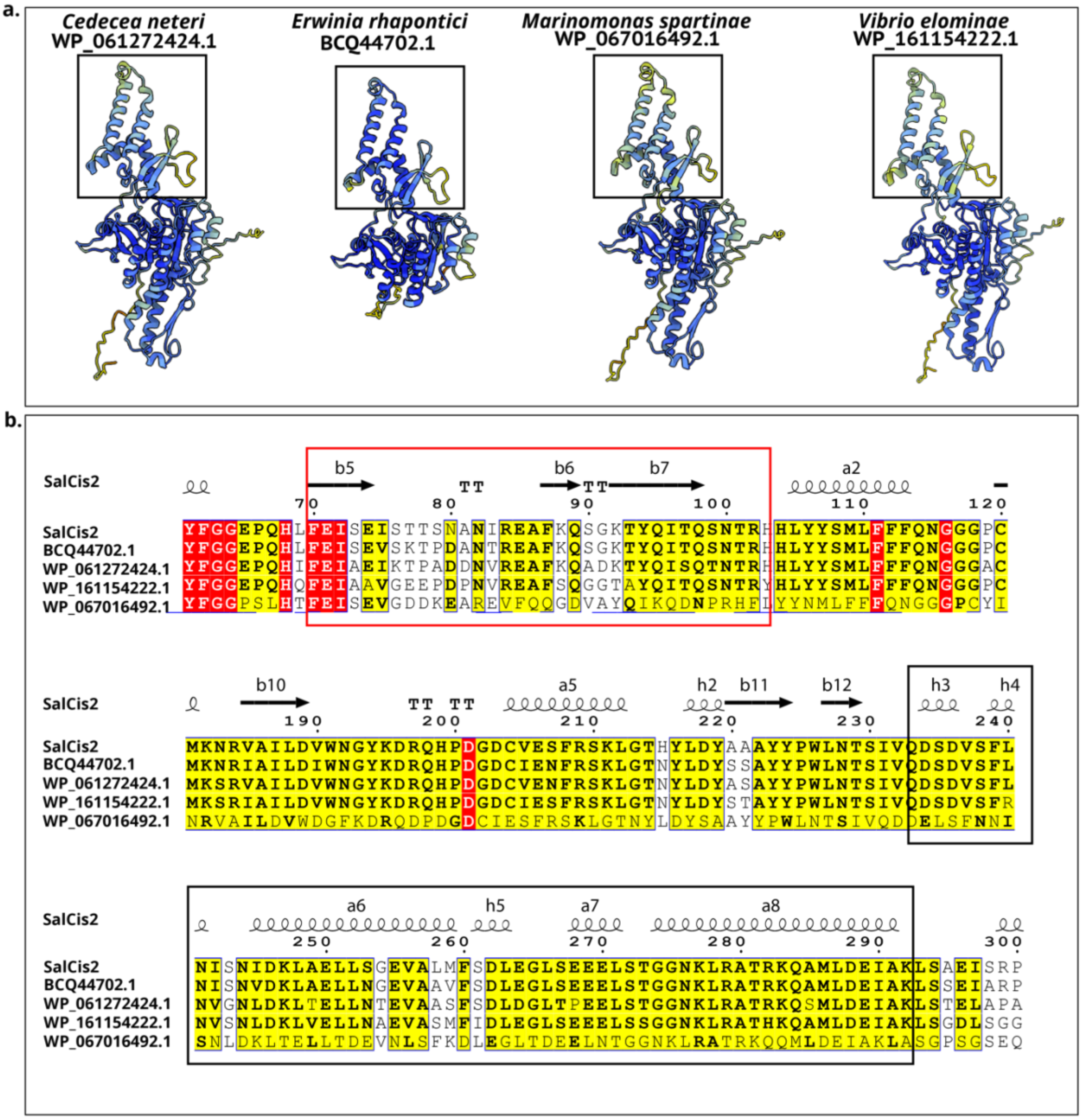


**Extended Data Figure 5. SalCis2 homologues, with unique features highlighted. predictions are coloured by pLDDT confidence score (blue to red from high to low).** **a:** Sheath protein predictions of SalCis2 homologues, black boxes indicate conserved vertical helix-loop-helix and X-loop motifs. **b:** Multiple sequence alignment of sheath proteins in b. Alignments truncated to highlight presence of X-loop. Red box highlights the X-loop motif, black boxes indicate the helix-loop-helix orientation motif

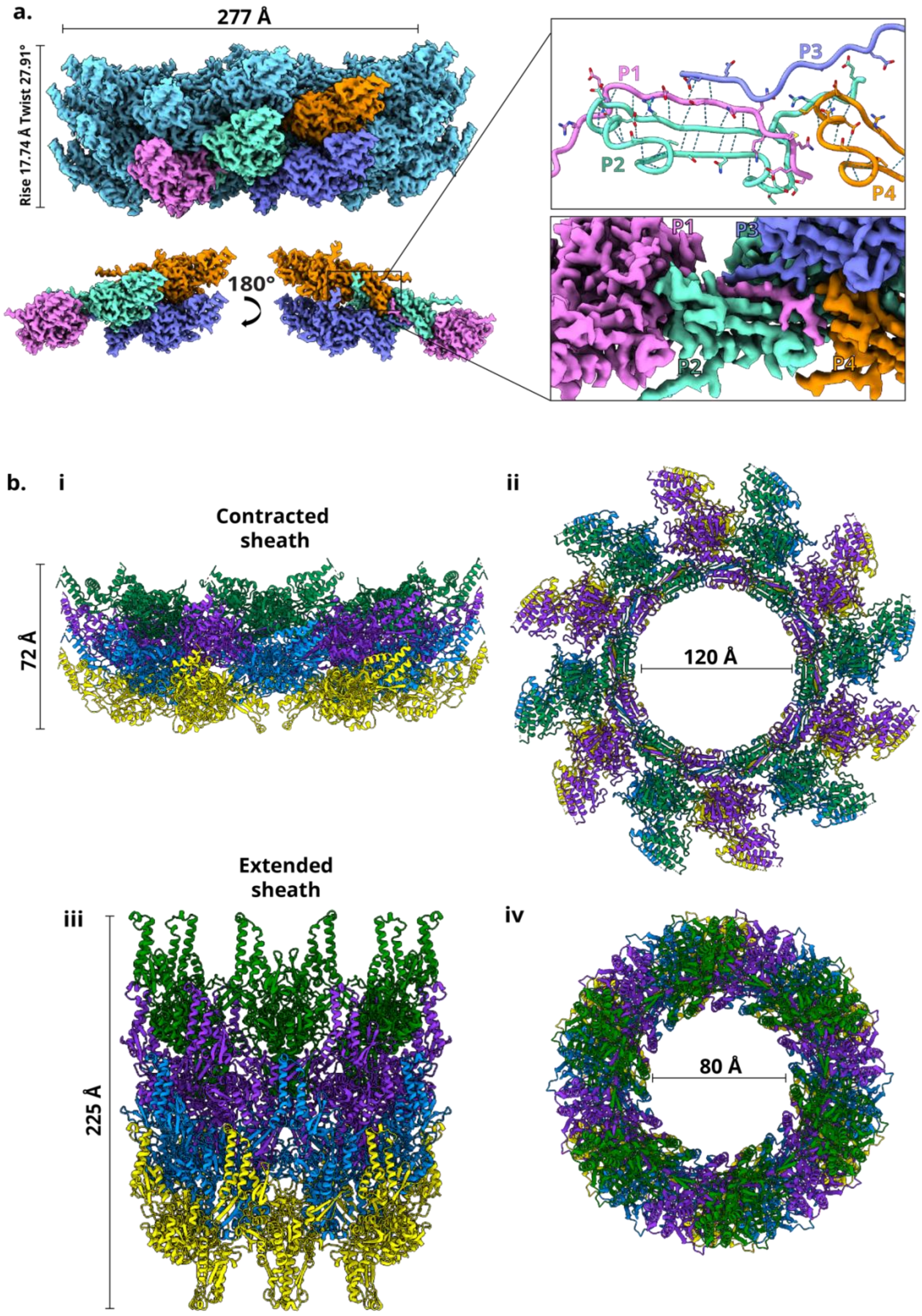


**Extended Data Figure 6. Cryo-EM structures of the SalCIS sheath.**

**a:** Cryo-EM reconstruction of the SalCIS sheath in the contracted state, side view depicts the arrangement of sheath protomers surrounding in contracted form, with individual protomers shown in distinct colours. Inset panels highlight the interface between four adjacent protomers (P1–P4), visualized in atomic detail (top) and corresponding cryo-EM density (bottom), revealing key inter-protomer contacts stabilizing the contracted conformation. **b:** Model of SalCIS sheath in the contracted (i-ii) and extended (ii and iv) conformations. 4 rings from each conformation are coloured according to ring. Structures are refined models expanded on to the cryoEM density. Inner tube is hidden in extended sheath model. Side and top views of each conformation show structural rearrangements accompanying sheath contraction.

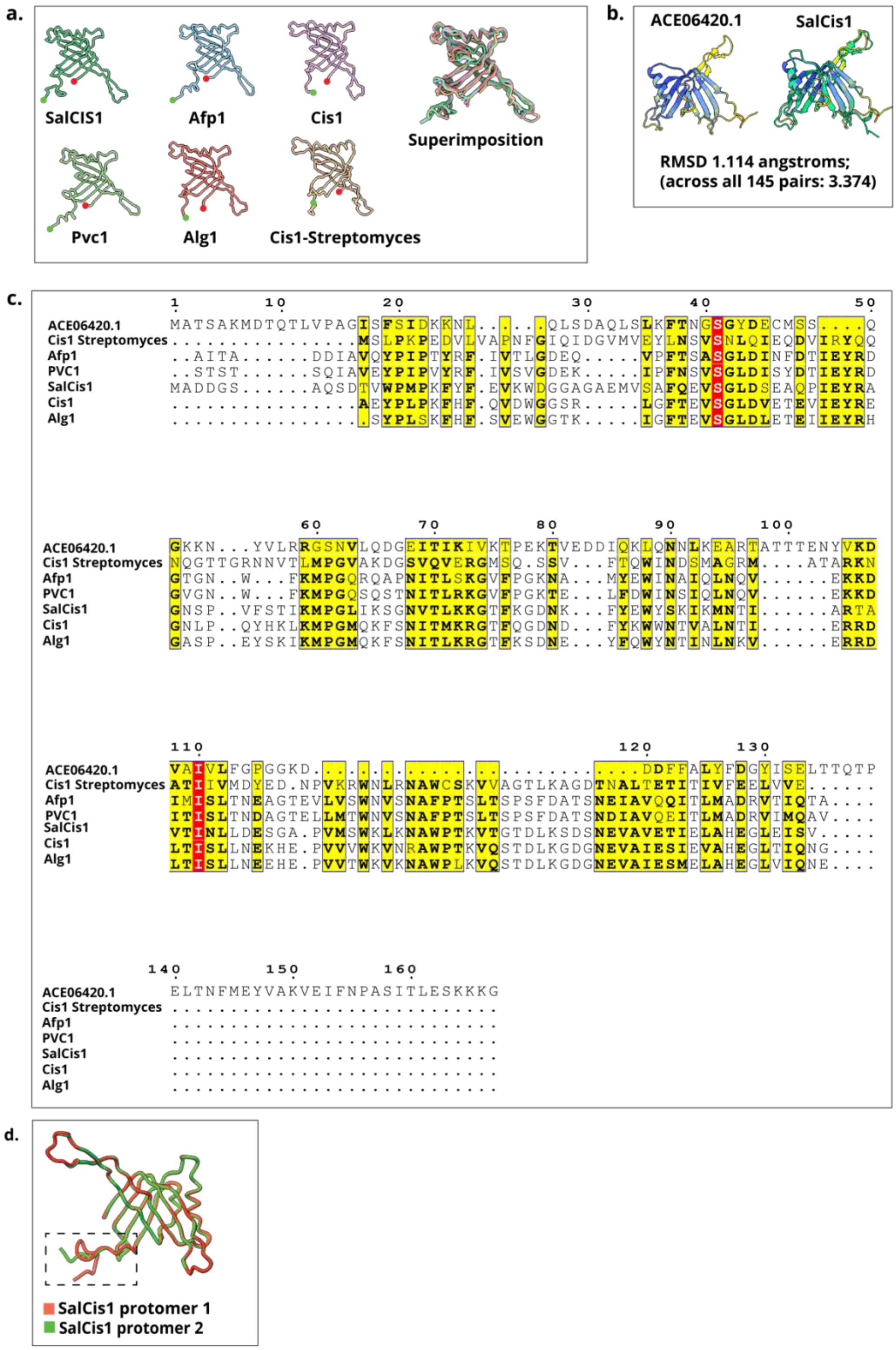


**Extended Data Figure 7. Comparison of SalCis1 to its inner tube homologues in other CIS, highlighting structural and sequence similarity**

**a:** Structural comparison of SalCis1 with inner tube proteins from other CIS systems, including superimposed structures highlight conserved structural similarity across systems. **b:** AlphaFold3 multimer predictions of protein homologue of SalCis1 in *Amoebophilus asiaticus*. **c:** Multiple sequence alignment of inner tube proteins in a. and b. **d:** Structural comparison of

SalCis1 protomers in the sheath (green) versus the cap (red). Black box highlights differences in C-terminal conformation

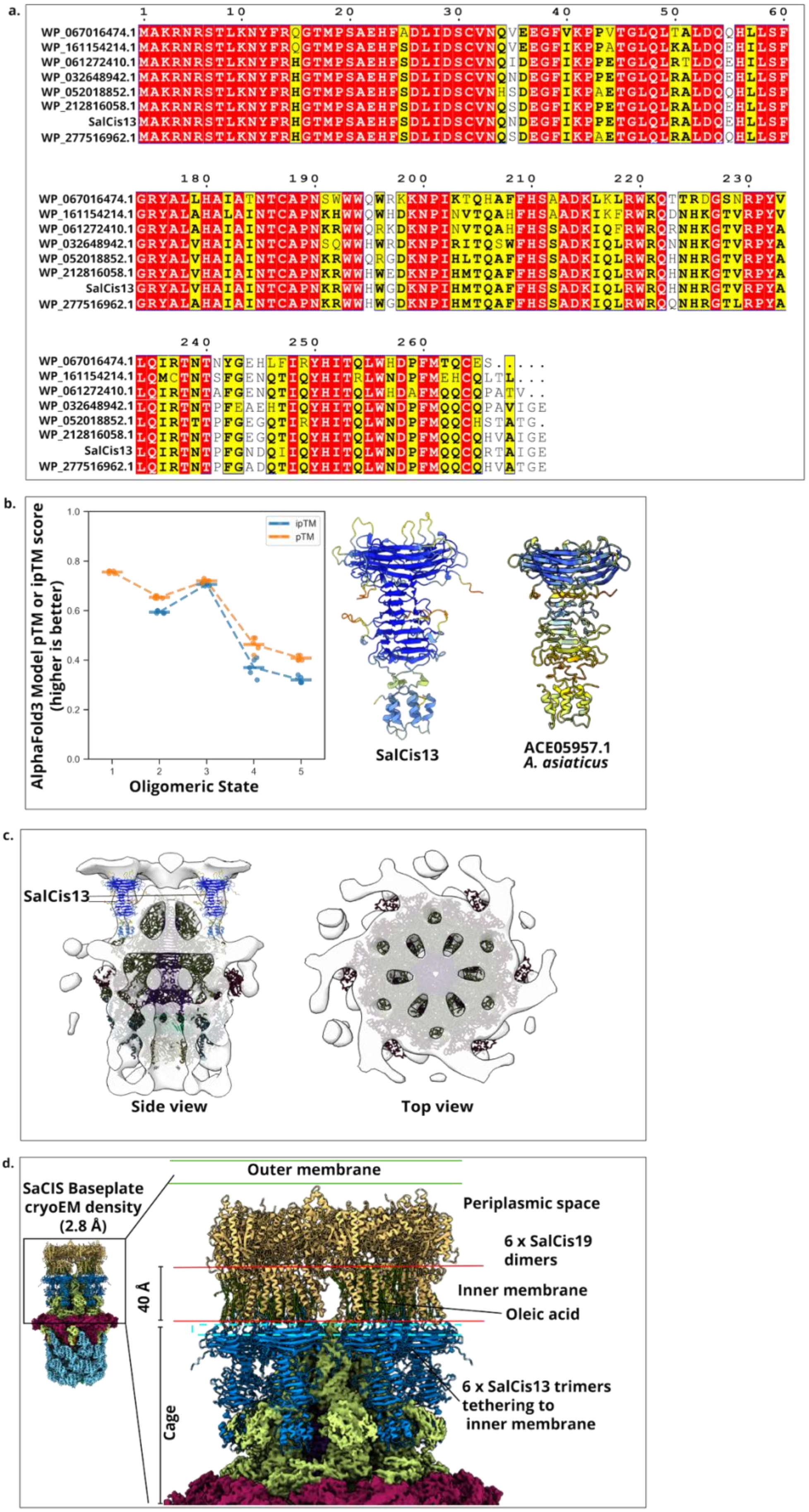

a.
WP_067016474.1
WP_161154214.1
WP_061272410.1
WP_032648942.1
WP_052018852.1
WP_212816058.1
SalCis13
WP_277516962.1
b.
AlphaFold3 Model pTM or ipTM score (higher is better)
ipTM
pTM
Oligomeric State
SalCis13
ACE05957.1
A. asiaticus
c.
SalCis13
Side view
Top view
d.
SaCIS Baseplate cryoEM density (2.8 Å)
Outer membrane
Periplasmic space
6 x SalCis19 dimers
Inner membrane
Oleic acid
40 Å
Cage
6 x SalCis13 trimers tethering to inner membrane

**Extended Data Figure 8. SalCis13 and its homologues resemble a membrane anchor**

**a:** MSA of SalCis13 homologues. Only N and C-terminus are shown, to highlight conserved similarity. Includes *Marinomonas spartinae*; WP_067016474.1, *Vibrio eleionamae*; WP_161154214.1, *Cedecea neteri*; WP_061272410.1, *Enterobacter*; WP_032648942.1, *Erwinia* sp.; WP_052018852.1, *Pectobacterium polaris*; WP_277516962.1. **b:** Left panel shows graph indicating favourable oligomeric state of SalCis13 is a trimer, when compared to a monomer, dimer, tetramer or pentamer. Right panel shows Alphafold3 prediction of SalCis13 as a trimer, alongside the same of the SalCis13 homologue in *A. asiaticus.* **c:** SalCIS baseplate complex and SalCis13 trimers fit into the sub-tomogram average of the *A. asiaticus* CIS to highlight structural and anchoring similarity. **d:** Proposed model of SalCis13 anchoring to the inner membrane via interaction with SalCis19. SalCis19 is predicted in complex with SalCis13 and SalCis11, and oleic acid to portray lipids in a membrane. Model was docked into a 2.8 Å reconstruction of the SalCIS baseplate. Blue lines demarcate probable location in periplasmic membrane, oleic acid-TMDs were measured at about 40 Å, similar to the average thickness of a lipid membrane. 6 predicted complexes were docked into the map, via SalCis11.

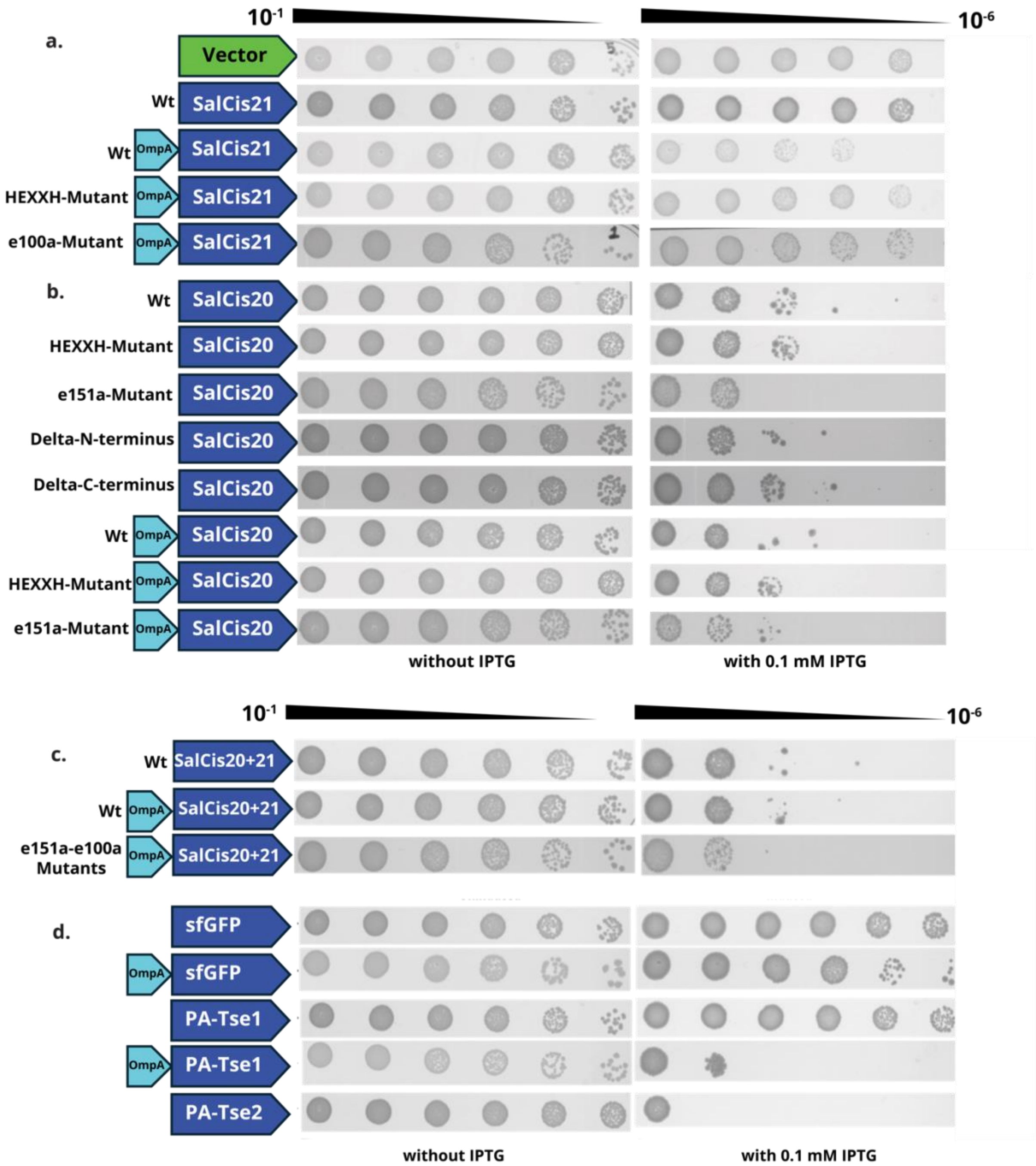


**Extended Data Figure 9. Toxicity spot assays of SalCIS toxins and controls. Cultures were serially diluted (10-fold) and spotted on agar plates with or without IPTG**
**a:** Spot dilution toxicity assay expressing SalCis21. **b:** Spot dilution toxicity assay expressing SalCis20. **c:** Spot dilution toxicity assay expressing SalCis20 and SalCis21 in tandem, fused with OmpA peptide, and point mutants fused with OmpA. **d:** Spot dilution toxicity assays of controls. sfGFP was used as an overexpression control. With and without OmpA peptide. **e:** *Pseudomonas aeruginosa* periplasmic toxin Tse1 without periplasmic *OmpA* sequence to show toxin overexpression control, fused to *OmpA* sequence to show rapid export into the periplasm and resulting toxicity. *Pseudomonas aeruginosa* cytosolic toxin Tse2 as a control to show cytosolic toxicity.
