## Supplementary Data for "Structure of a contractile injection system in *Salmonella enterica* subsp. *salamae*"

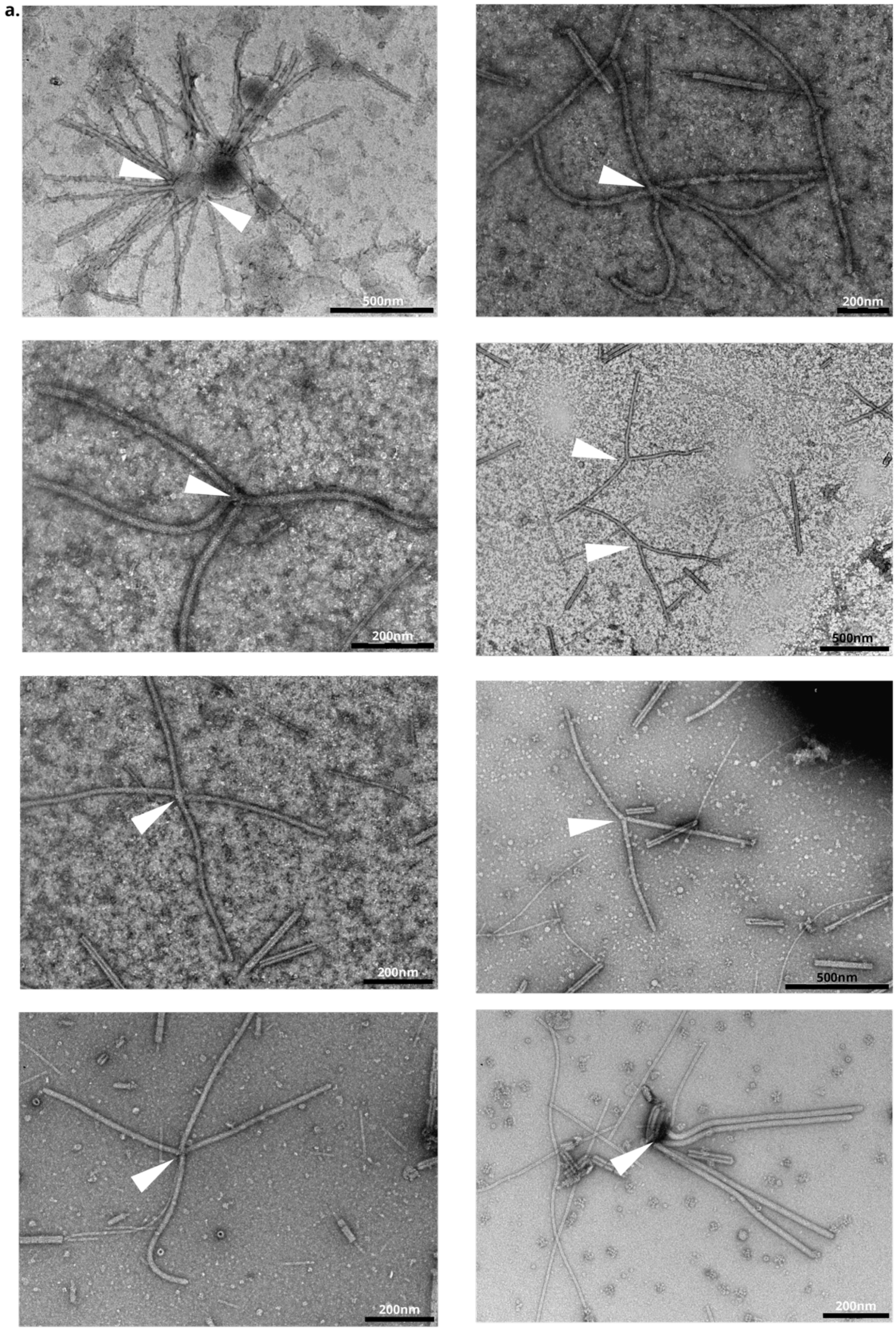


**Supplementary Figure 1. Negative-stain electron micrographs of *S. enterica* eCIS particles forming bundles with embedded baseplates**

**a:** Representative negative-stain transmission electron microscopy (TEM) images of *Salmonella enterica* particles, visualized using a FEI Morgagni TEM at 80kV. Particles are observed assembling into bundles, with baseplates embedded in some densities. White arrows indicate baseplates positioned within these bundled structures. Here the entire SalCIS operon is expressed in *E. coli.*

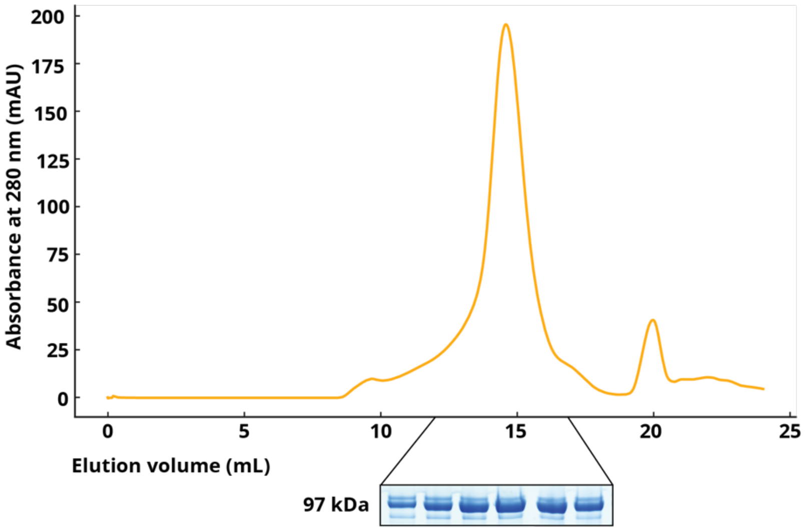


**Supplementary Figure 2. SEC profile of purified SalCis19.** SEC was performed on a Superdex 200 GL column, GE Healthcare, equilibrated with running buffer and 0.02% LMNG. UV absorbance trace at 280 nm is shown. Coomassie-stained SDS-PAGE of selected fractions is displayed, showing the presence of SalCis19 (97kDa). The *E. coli* protein SlyD was detected using mass-spec in the band below SalCis19 but was not present in the cryoEM dataset.

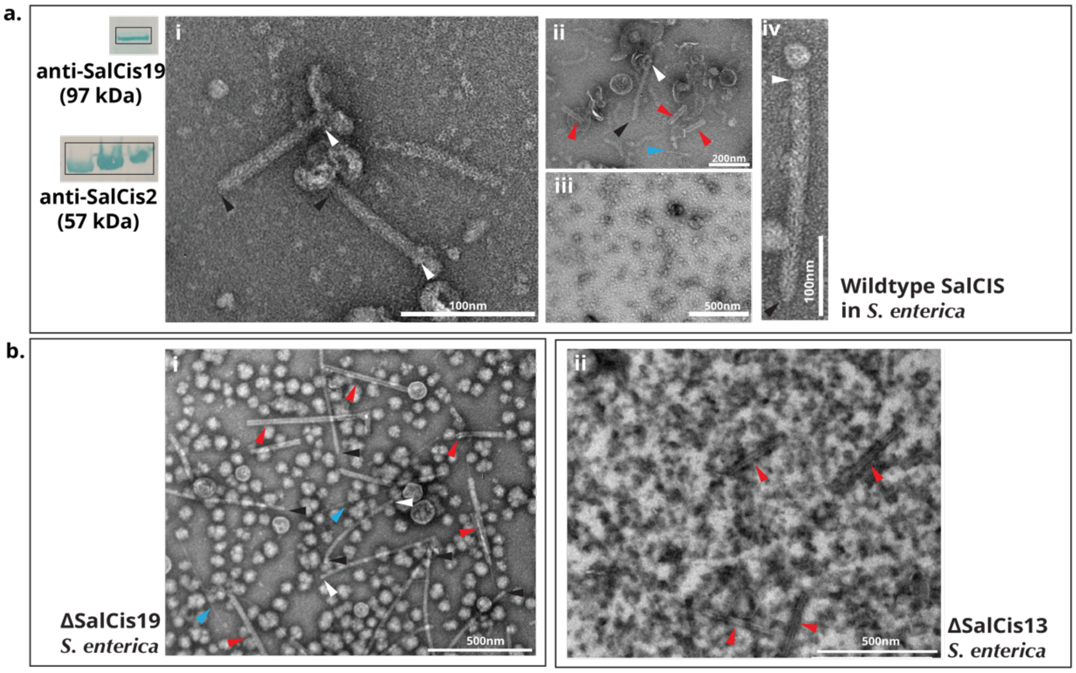


**Supplementary Figure 3. Negative-stain TEM images of SalCIS.**
Black arrowheads denote caps, white arrows denote baseplates, red arrows deonote contracted sheath assemblies, blue arrows denote expelled inner tubes. Western blot bands are cropped for clarity. All uncropped Western blot images are available as a separate file. **a:** (i) Low magnification image of diluted final preparation from fractions 6, 7 and 8, after sucrose gradient fractionation, of endogenously expressed wild-type SalCIS in *S. enterica* subspecies *salamae*, resulting in a crude preparation with very few assembled particles. (ii)

Second image of fractions 6, 7 and 8, showing mostly contracted sheaths (iii) TEM image of fraction 2, and Western blot of the same, band indicates presence of SalCis19 at 97 kDa, probed with an anti-SalCis19 specific antibody. Western blot band seen at 57 kDa in a blot of the same fractions, as a positive control that SalCIS is expressed in the *S. enterica* subspecies *salamae*. (iv) Magnified view of one extended SalCIS particle from the purification. **b:** (i) Low magnification image of diluted final preparation from fraction 8 and 9, after iodixanol gradient fractionation, of heterologously expressed SalCIS ΔSalCis19 mutant in *E. coli* (BL21 DE3), resulting in a crude preparation. (ii) Low magnification image of diluted final preparation from fraction 8 and 9, after iodixanol gradient fractionation, of heterologously expressed SalCIS ΔSalCis13 mutant in *E. coli* (BL21 DE3).

**a.**

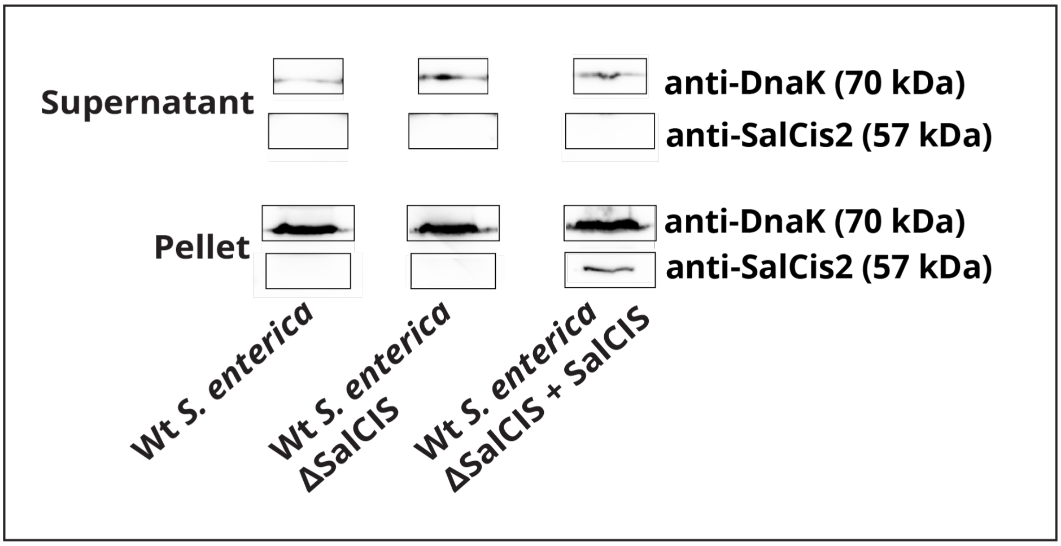


**b.**

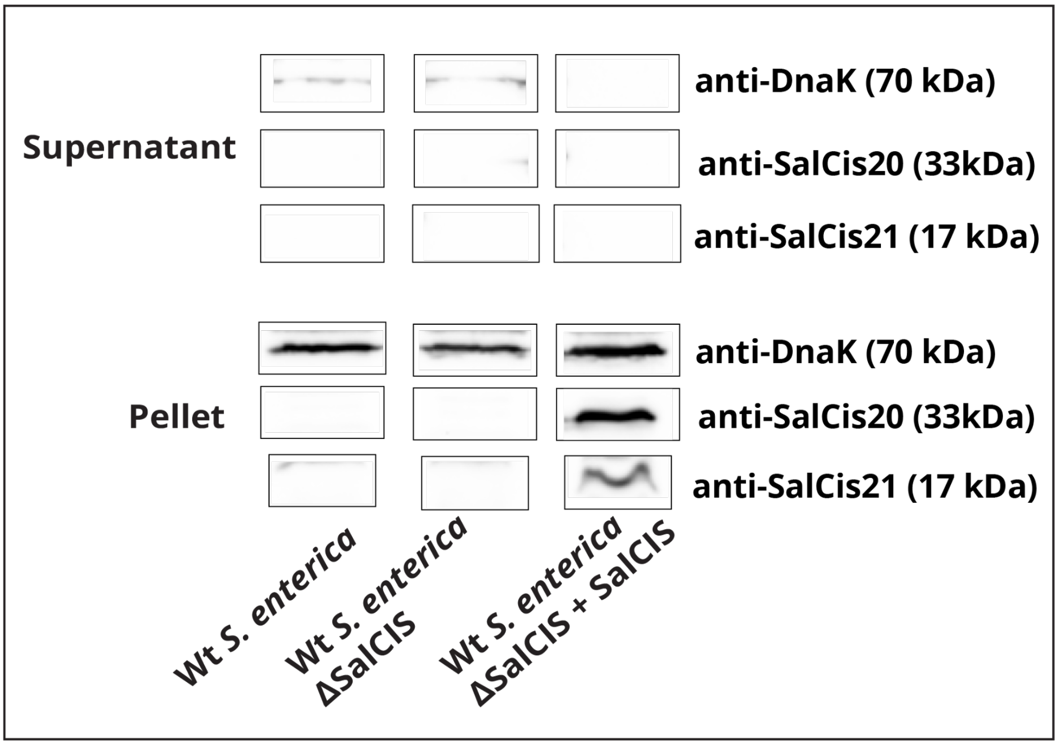

**Supplementary Figure 4. Western blot analysis of secretion assays.** Culture supernatants and cell pellets were separated by SDS-PAGE and probed with antibodies against SalCis2 (a.), SalCis20(c.) and SalCis21 (c.). Absence of proteins in the supernatant indicates no secretion. Loading controls for whole-cell fractions are shown using anti-DnaK antibody. Bands are cropped for clarity.

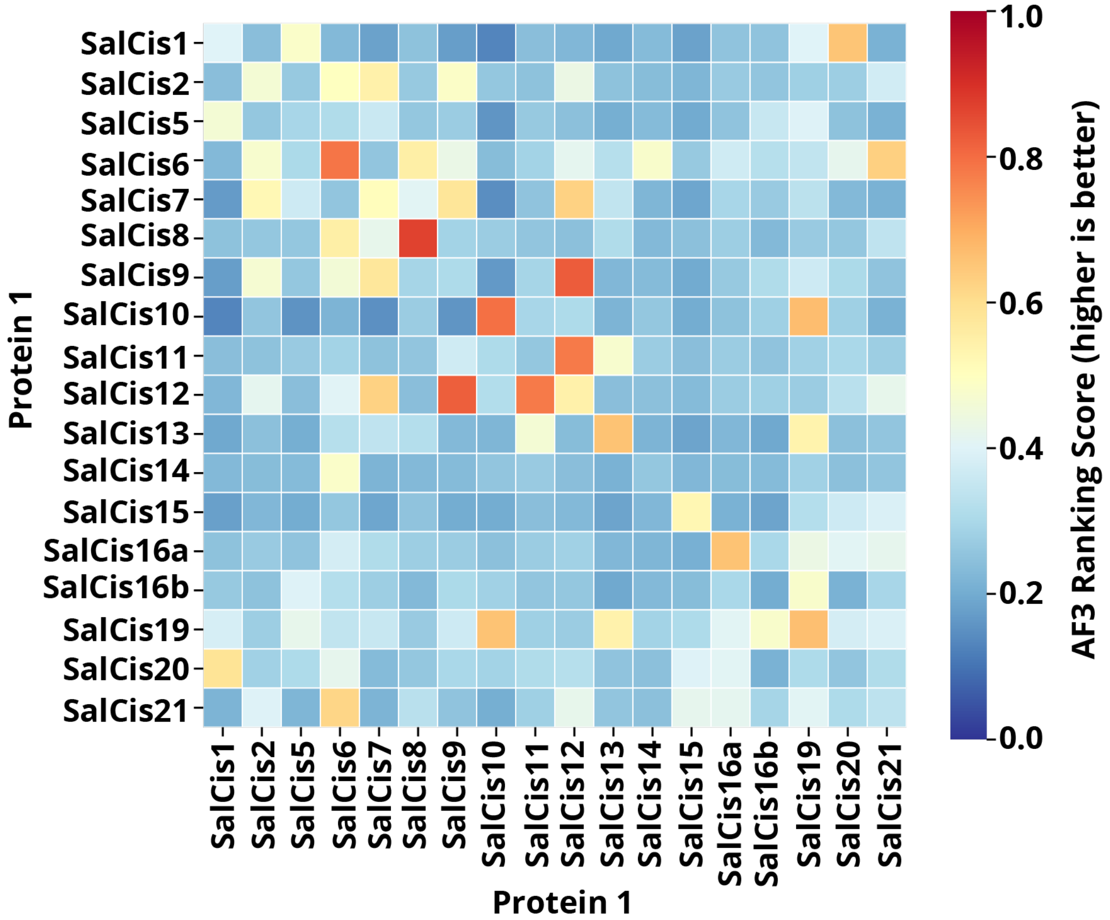


**Supplementary Figure 5. Heat map of AlphaFold3 predicted protein interactions of all proteins in the *S. enterica* CIS operon versus each other.** Higher the ranking, more likely the interaction. AlphaFold multimer 3 was used to predict all interactions. For the all-vs-all analysis, 5 models were created at 5 random seeds, creating 25 models for analysis. The value shown in the heatmap is the mean average of the scores calculated for the 25 models.

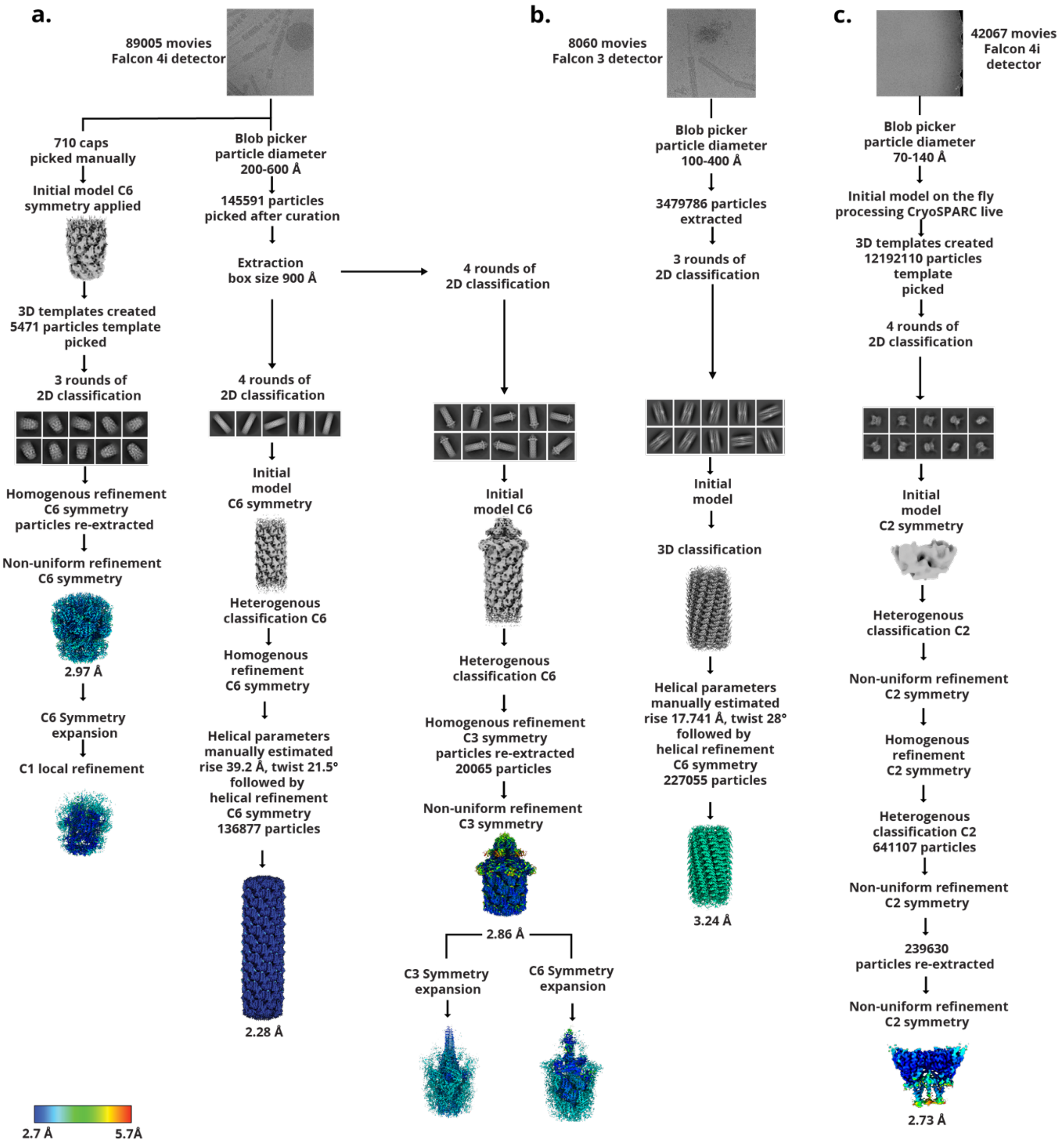


**Supplementary Figure 6. Cryo-EM workflow.**

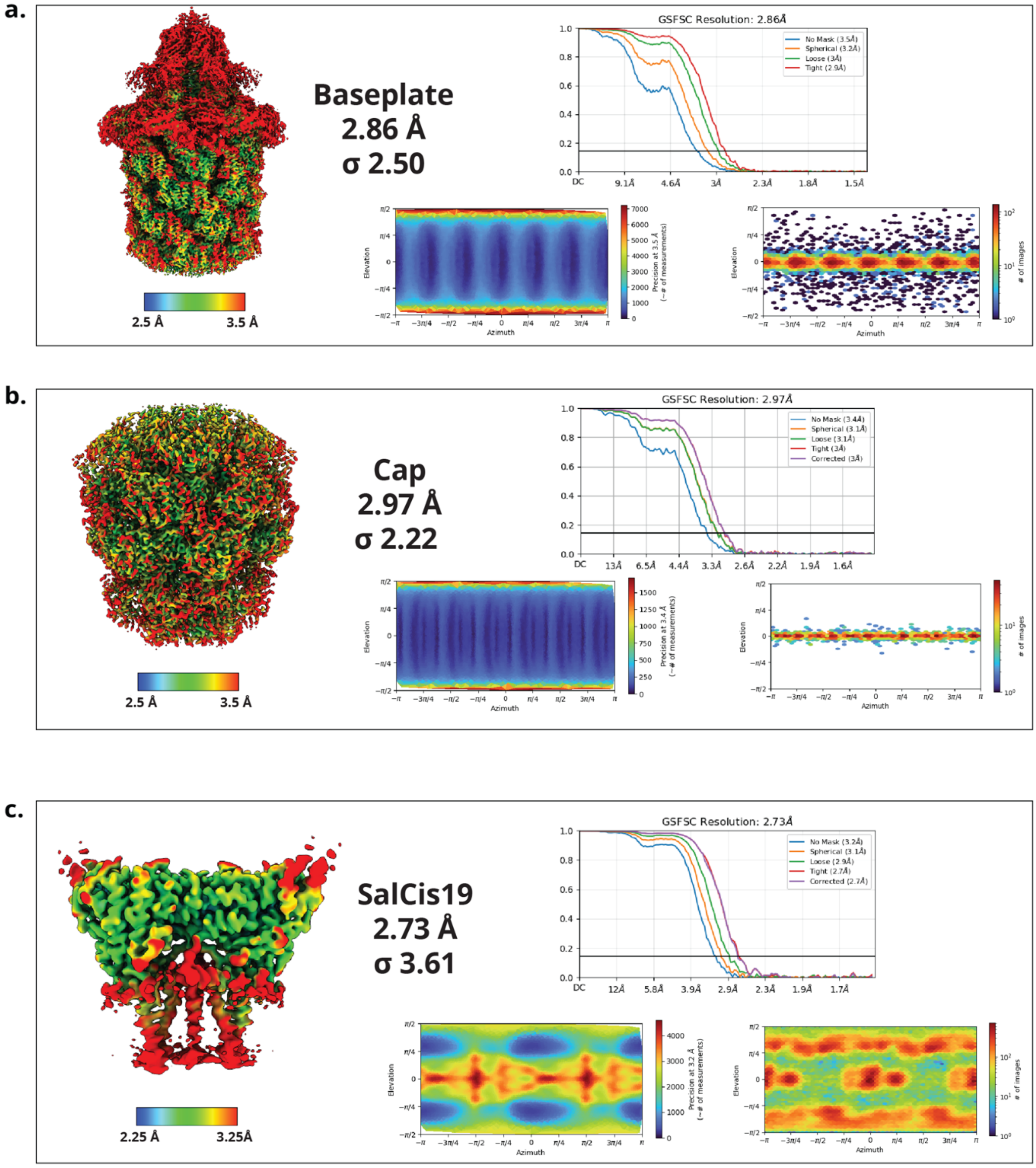


**Supplementary Figure 7. Masked FSC curves**. Local resolution maps (left), masked FSC curve (right, top), direction distribution (bottom left) and view direction (bottom right) of baseplate (a), cap (b) and SalCis19 (c).

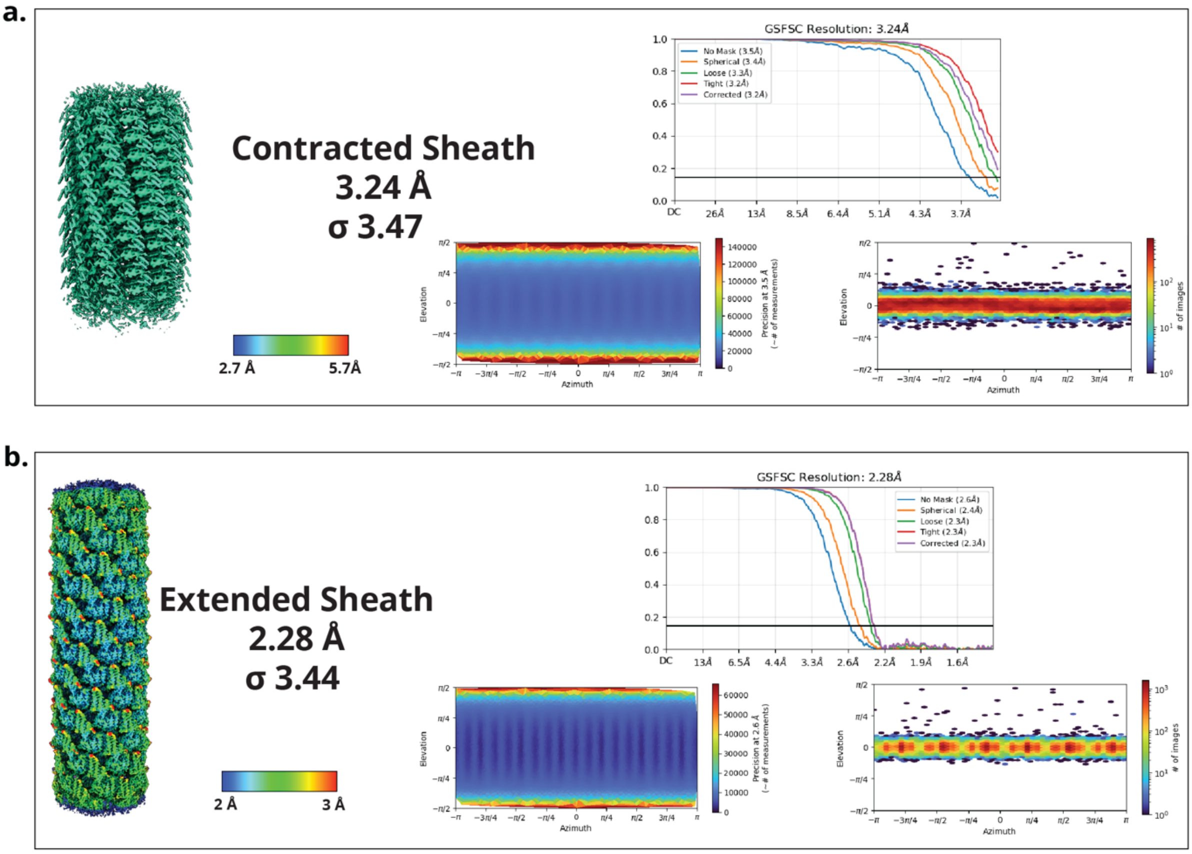


**Supplementary Figure 8. Masked FSC curves.** Local resolution maps (left), masked FSC curve (right, top), direction distribution (bottom left) and view direction (bottom right) of contracted sheath (a) and extended sheath (b).

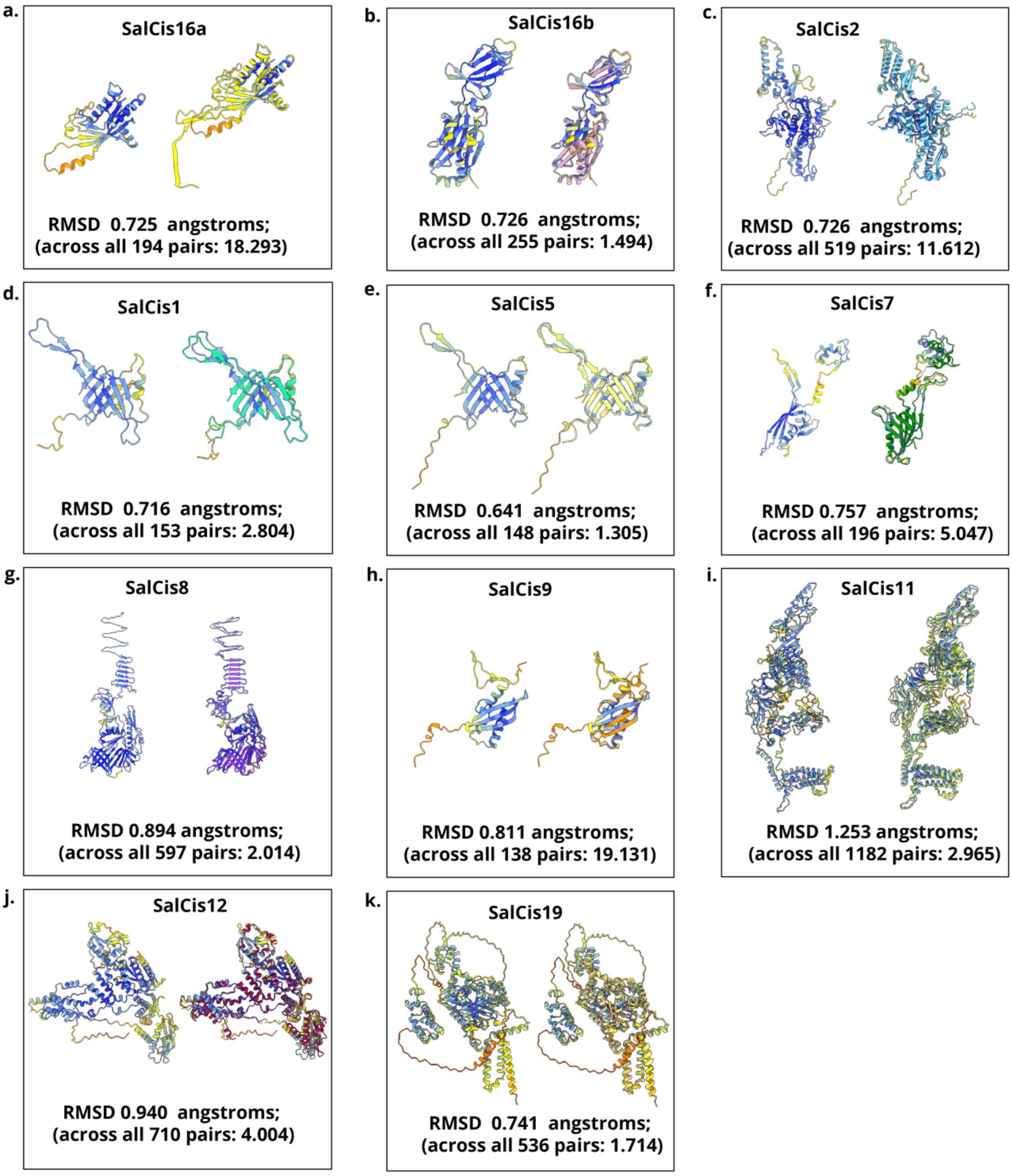


**Supplementary Figure 9. Comparison of experimentally determined SalCIS protein structures with AlphaFold3 predictions.**

a-k. For each protein, the AlphaFold3-predicted structure (left in each panel) is shown superimposed with the experimentally determined structure (right in each panel). Models are coloured by chain, while AlphaFold3 predictions are coloured by pLDDT confidence score (blue to red from high to low).

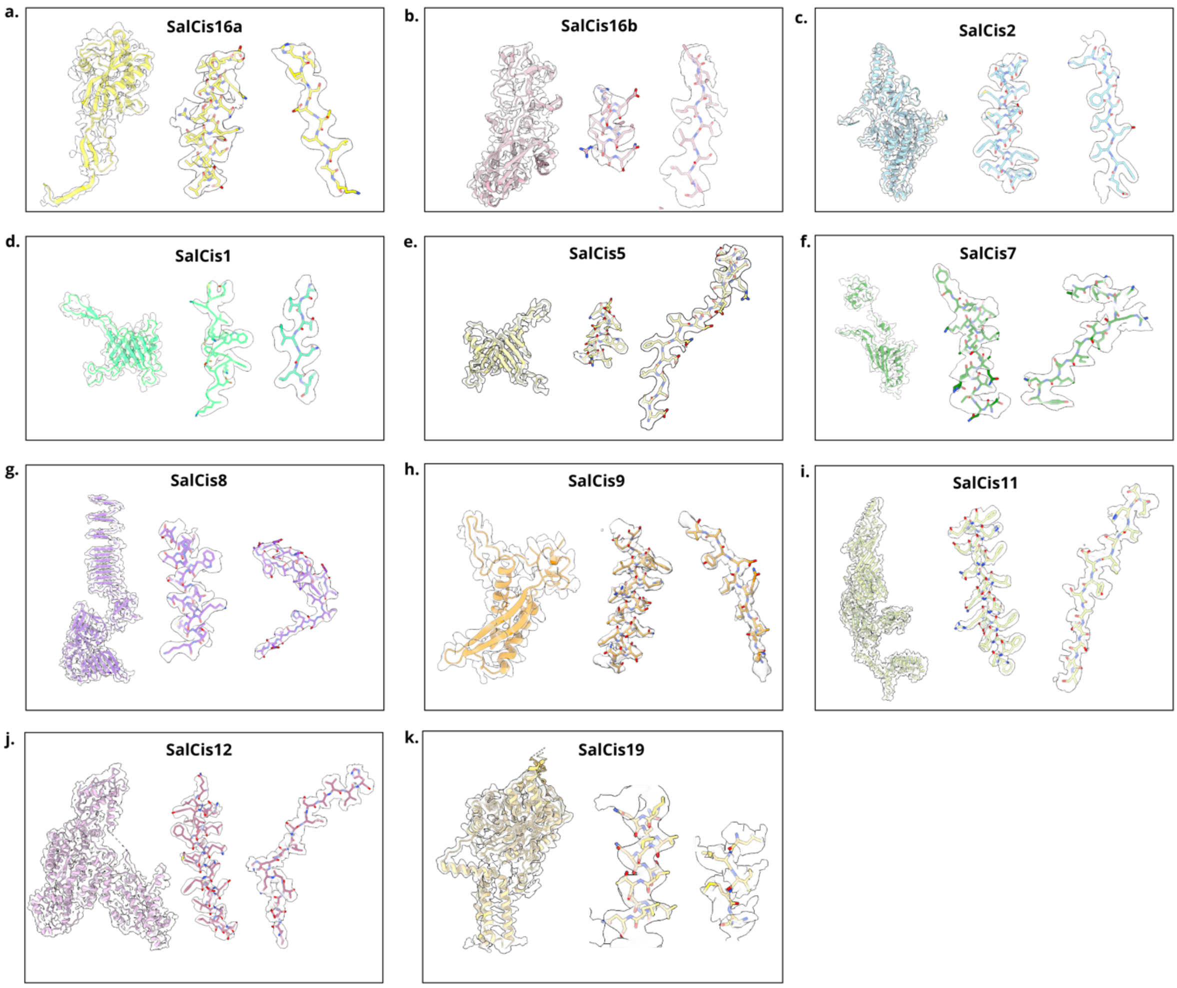


**Supplementary Figure 10. Modelled SalCIS proteins within cryo-EM density maps.**
a-k. Cryo-EM models of all SalCIS components shown within their corresponding cryo-EM densities, segmented, opacity set to 30% (left), alongside magnified views of a representative α-helix (middle) and β-strand (right) region.

**Supplementary Formula 1.**

**Contraction length estimate:**
SalCIS has an estimated 918 sheath subunits. The extended length of the system was approximated at 600 nm. Using the helical parameters for the extended and contracted sheath structures: extended rise, 39.2 Å; contracted rise: 17.7 Å. Therefore, the ratio between the two lengths is (39.2 Å / 17.7 Å) = 2.2. Using this ratio, we can approximate the length of the contracted state to be: 600 nm / 2.2 = 272 nm, which means the contraction distance is (600 nm - 272 nm) 328 nm.

**Supplementary Table 1.** Summary of structural modelling, sequence coverage, and relative abundance of SalCIS proteins. Residues modelled and not modelled for each SalCIS protein are indicated based on cryo-EM density map interpretations. Map type refers to the refinement strategy applied, including local refinements with C1 symmetry and symmetry-expanded reconstructions or helical refinement where applicable. LC-MS sequence coverage represents the percentage of the entire operon sequence identified from purified sample. Relative abundance was determined from LC-MS analysis and normalized to the most abundant component, SalCis16b. Dashes indicate proteins for which no structural model was built or data was unavailable.

| Protein | Residues Modelled | Residues Not Modelled | Map Type | LCMS sequence coverage of entire operon expressed and purified % | Relative abundance | MS/MS count |
|---|---|---|---|---|---|---|
| **SalCis16a** | 1-62,71-193 | 63-70,194 | Local, C1 refinement, symmetry expanded C6 | 83.5 | 0.257 | 380 |
| **SalCis16b** | 3-124,136-257 | 1-2,125-135,258-260 | Local, C1 refinement, symmetry expanded C6 | 97.3 | 1.000 | 1,589 |
| **SalCis2-extended** | SalCis2_R1:3-516<br>SalCis2_R2:3-516<br>SalCis2_R3:3-515 | SalCis2_R1:1-2,517-519<br>SalCis2_R2: 1-2,517-519<br>SalCis2_R3: 1-5,516-519 | Helical refinement, symmetry C6 | 90.8 | 0.753 | 4,185 |
| **SalCis2-contracted** | 1-264,278-519 | 265-277 | Helical refinement, symmetry C6 | - | - | - |
| **SalCis1** | 4-156 | 1,2,3 | Local, C1 refinement, symmetry expanded C3 | 99.4 | 0.501 | 931 |
| **SalCis5** | 11-41,43,45-49,51-77,82-100,102-111,113-158 | 1-10,42,44,50,101,112 | Local, C1 refinement, symmetry expanded C3 | 86.7 | 0.035 | 57 |
| **SalCis6** | 49-56 | 1-48 | Local, C1 refinement, symmetry expanded C3 | 98.2 | 0.026 | 73 |
| **SalCis7** | 1-159,167-211 | 160-166,212-213 | Local, C1 refinement, symmetry expanded C3 | 98.2 | 0.163 | 613 |
| **SalCis8** | 3-597 | 1-2 | Local, C1 refinement, symmetry expanded C3 | 79.1 | 0.040 | 583 |
| **SalCis9** | 6-143 | 1-5,144-149 | Local, C1 refinement, symmetry expanded C6 | 83.2 | 0.009 | 59 |
| **SalCis11** | 2-34,36,40-50,55-137,140-141,144-155,175,177-296,298-315,331-397,399-400,419-420,422-433,436-438,441-444,450-452,454-457,463- | 1,37-39,51-54,138-139,142-143,156-174,176,297,316-330,398,401-418,421,434- | Local, C1 refinement, symmetry expanded C6 | 66.6 | 0.016 | 706 |

| | | | | | | |
|---|---|---|---|---|---|---|
| | 484,497-513, 515-531,534-535,540-541,543-549,555-558,560-585,592-773,776-812,815-862,867-885,893-955,957-1128,1135-1237,1241-1247,1252-1257,1262-1276,1278-1285 | 435,439-440,445-448,449,453,458-459,460-462,485-495,496,514,532-533,536-539,542,550-554,559,586-591,774-775,813,814,863-866,886-891,892,956,1129-1134,1238-1240,1248-1251,1258-1261,1286-1287,1288-1323,1277 | | | | |
| **SalCis12** | 15-133,137-152,170-265,276-327,336-535,546-574,577-584,588-589,595-607,609-614,616-624,626-627,642-643,652-658,662-683,685-798 | 1-14,134-136,153-169,266-275,328-335,536-545,575-576,585-587,590-594,608,615,625,628-641,644-651,659-661,684,799-804 | Local, C1 refinement, symmetry expanded C6 | 72.9 | 0.021 | 472 |
| **SalCis19** | 347,349,351-353,355-541,543-602,604-633,635-799,801-842,844-848,850,853-858,860 | 348,350,354,542,603,634,800,843,849,851-852,859,861-864 | C2 | 50.8 | 0.004 | 257 |
| **SalCis13** | - | - | - | 54.3 | 0.005 | 70 |
| **SalCis15** | - | - | - | 86.7 | 0.406 | 1,327 |
| **SalCis20** | - | - | - | 86.2 | 0.294 | 701 |
| **SalCis21** | - | - | - | 67.1 | 0.042 | 230 |

**Supplementary Table 2.** SalCIS proteins and their homologues. HHPred was used inside the MPI Toolkit, to search for protein sequence based structural homology of all proteins in the cluster[49]. AlphaFold3-Multimer was used to predict structures of all proteins in the CIS cluster, for all four *S. enterica* strains[50]. The predicted structures were then submitted to the DALI server (Ekhidna 2 biocenter) to survey for structurally homologous proteins[51]

| SalCis Protein | Homologous Proteins |
|---|---|
| **SalCis1** | Hcp (T6SS); Afp1 (Afp); Pvc1 (Pvc); Alg1 (AlgoCIS); Cis1 (*Streptomyces*) |
| **SalCis2** | Sheath proteins: Afp2 (Afp), Pvc2 (Pvc), Cis2 (Streptomyces), Alg2 (AlgoCIS), ACE06419.1 (A. asiaticus) |
| **SalCis5** | Tube-related proteins: Alg5 (AlgoCIS), Pvc5 (Pvc), Afp5 (Afp), gp19 |
| **SalCis6** | Plug protein; structurally similar to SalCis8 loop region |

| | |
|---|---|
| **SalCis7** | Baseplate wedge proteins: Alg7 (AlgoCIS), Cis7 (*Streptomyces*), ACE06424.1 (*A. asiaticus*) |
| **SalCis8** | Spike complex: gp5/gp27 (T4 phage); PAAR-like proteins |
| **SalCis9** | Sheath initiator |
| **SalCis10** | PAAR-domain proteins; T6SS VgrG complex; phiKZ gp163.1 |
| **SalCis11** | Baseplate cage: gp6 (T4 phage), Alg11 (AlgoCIS), Cis11 (*Streptomyces*), ACE05958.1 (*A. asiaticus*) |
| **SalCis12** | Baseplate adapter: gp7 (T4 phage), Alg12 (AlgoCIS), Cis12 (*Streptomyces*), ACE06428.1 (*A. asiaticus*) |
| **SalCis13** | Adhesin-like proteins: Bd2133 (MAT domain); ACE05957.1 (*A. asiaticus*) |
| **SalCis15** | ATPase: AAA+ ATPases (Afp15, Pvc15) |
| **SalCis16a** | Cap protein: Pvc16 (Pvc), Afp16 (Afp), Cis16 (*Streptomyces*) |
| **SalCis16b** | Cap-related protein: Alg16b (AlgoCIS), ACE06418.1 (*A. asiaticus*) |
| **SalCis19** | Peptidoglycan hydrolase: SleL (*Bacillus cereus*) |
| **SalCis20** | T6SS-associated toxin: DUF4157-domain toxin |
| **SalCis21** | Toxin: VgrG2b C-terminal toxin (*Pseudomonas aeruginosa*), HEXXH metalloprotease domain |

### **Supplementary Table 3.** CryoEM data collection parameters and validation statistics

| | **Baseplate** | **Extended Sheath** | **Cap** | **SalCis19** | **Contracted Sheath** |
|---|---|---|---|---|---|
| **Data collection and processing** | | | | | |
| Microscope | Titan Krios G2 | | | | Glacios |
| Voltage (kV) | 300 | | | | 200 |
| Magnification (nominal) | 96,000x | | | | 73,000x |
| Detector | Falcon 4i | | | | Falcon 3 |
| Pixel size (Å) | 0.73 | | | | 1.6 |
| Defocus range (μm) | -0.5 to -2.5 | | | | -0.5 to -2.5 |
| Frames/movie | 41 | | | | 40 |
| Total exposure dose ($e^-/Å^2$) | 42 | | | | 36.84 |
| Movies | 80000 | | | 42067 | 8060 |
| Number of particles (initial) | 145591 | 145591 | 5471 | 12192110 | 3479786 |
| Number of particles (final) | 20065 | 136877 | 5471 | 239630 | 227055 |
| Box size (pixels) | 630 | 900 | 360 | 320 | 400 |
| Symmetry imposed | C3 | C6 | C6 | C2 | C6 |
| Map resolution (Å) (FSC 0.143) | 2.86 | 2.28 | 2.97 | 2.73 | 3.24 |
| **Refinement** | | | | | |
| Refinement type | 1 C3 asymmetric unit | Helical | 1 C6 asymmetric unit | 1 Dimer | Helical |
| SalCIS entities | SalCis11, SalCis12, SalCis9, SalCis2, SalCis8, SalCis10, SalCis6 | SalCis2, SalCis1 | SalCis16a, SalCis16b, SalCis2, SalCis1 | SalCis19 | SalCis2 |
| Non-hydrogen atoms | 99395 | 184968 | 17243 | 17311 | 137826 |
| Protein residues | 6276 | 12006 | 1102 | 1071 | 8910 |
| RMSZ Bond lengths | 0.003 | 0.035 | 0.003 | 0.002 | 0.002 |
| RMSZ Bond angles | 0.538 | 0.692 | 0.557 | 0.439 | 0.444 |
| CC (mask) | 0.79 | 0.9 | 0.8 | 0.82 | 0.89 |
| CC (volume) | 0.78 | 0.89 | 0.8 | 0.82 | 0.84 |
| CC (peaks) | 0.01 | 0.37 | -0.16 | 0.54 | 0.35 |
| B-factors (mean;Å²) | 101.79 | 84.88 | 105.22 | 137.26 | 47.48 |

| | | | | | |
|---|---|---|---|---|---|
| Refinement resolution (FSC map vs model (masked = 0.143) Å | 2.77 | 2.25 | 2.95 | 2.6 | 3.19 |
| **Validation** | | | | | |
| MolProbity score | 1.21 | 1.26 | 1.06 | 1.12 | 0.88 |
| Clash score, all atoms | 2.31 | 4.93 | 1.97 | 3.18 | 1.45 |
| Rotamer outliers (%) | 1.33 | 0.41 | 1.04 | 1.04 | 0.28 |
| Ramachandran Favoured (%) | 97.46 | 98.68 | 97.62 | 98.4 | 98.88 |
| Ramachandran Allowed (%) | 2.51 | 1.22 | 2.29 | 1.6 | 1.12 |
| Ramachandran Outliers (%) | 0.03 | 0.1 | 0.09 | 0 | 0 |
